# Sequential accumulation of adaptive alleles forms an inversion supergene in deer mice

**DOI:** 10.64898/2026.09.23.753864

**Authors:** Olivia S. Harringmeyer, Shuonan He, Hopi E. Hoekstra

**Author notes:** These authors contributed equally (Olivia S. Harringmeyer, Shuonan He).

## Abstract

Supergenes are clusters of co-inherited loci that affect multiple or complex phenotypes. Despite the growing number of chromosomal inversions identified as supergenes in natural populations, their molecular basis and evolutionary history often remain obscure. Here, we identified two candidate genes, *Slc45a2* and *Npr3*, within a 41-Mb inversion supergene in the deer mouse (*Peromyscus maniculatus*) that respectively drive darker coats and longer tails—two traits associated with forest adaptation. Mice homozygous for the inversion (*inv/inv*) exhibit elevated *Slc45a2* expression in melanocytes relative to the congenic standard genotype (*std/std*), disrupting pheomelanin production. In parallel, downregulation of *Npr3* in *inv/inv* mouse growth plates prolongs postnatal growth of caudal vertebrae, resulting in tail elongation. Population-level analyses further implicate that this supergene arose through the subsequent accumulation of the *Npr3* allele within the inversion, rather than by capturing all beneficial mutations at its origin.

## Main Text

Adaptation is a complex process that involves coordinated changes in multiple traits. How such changes are encoded and maintained at the genetic level remains a long-standing question. Supergenes––genomic regions comprising tightly linked loci that segregate as single Mendelian units––drive multi-trait phenotypes and serve as a quintessential mechanism that facilitates local adaptation across diverse taxa (*1–3*).

Chromosomal inversions offer an ideal structural foundation for supergenes, as they suppress recombination with the ancestral (herein referred to as the “standard”) haplotype, thereby preserving the linkage of adaptive alleles (*4–7*). Empirical studies demonstrate that inversion supergenes are widespread, orchestrating phenotypes ranging from wing pattern mimicry in the *Heliconius* butterflies to diverse mating strategies in the ruff (*8–11*). Despite these compelling examples, a fundamental question persists: how does an inversion become a supergene? The “capture model” posits a mechanism by which an inversion can immediately function as a supergene, when it traps multiple preexisting adaptive alleles and/or influences gene function through its breakpoints, thus conferring an instantaneous selective advantage over the standard haplotype (*1, 4*). In contrast, an inversion may arise first and only subsequently acquire additional adaptive mutations which transform it into a supergene over time, hereafter referred to as the “accumulation model” (*1, 12*). To date, there is limited experimental evidence in support of either model. Distinguishing between them requires not only a mechanistic dissection of the supergene, but also a reconstruction of its evolutionary history (*1, 7*).

The deer mouse (*Peromyscus maniculatus*), the most abundant native mammal in North America (*13, 14*), displays remarkable inversion polymorphisms among its natural populations, making it a powerful model for studying supergene evolution (*15*). One such inversion supergene has been identified using two adjacent deer mouse populations in the northwestern US, where the forest ecotype (*P.m.rubidus*) possess significantly darker coats and longer tails compared to their prairie-dwelling conspecifics (*P.m.gambelii*) (Fig. 1A) (*16*). Forward genetic mapping linked both traits to a 41-Mb inversion on chromosome 15 (Fig. 1B), whose allele frequency sharply declines along the forest-prairie habitat transition, consistent with local adaptation (*16*). Given its estimated large effect sizes on tissue-specific phenotypes (explaining 46% of coat color variation and 12% of tail length variation), this inversion presents an ideal opportunity to understand the molecular mechanisms and evolutionary trajectory that gave rise to a supergene.

**Figure 1.**
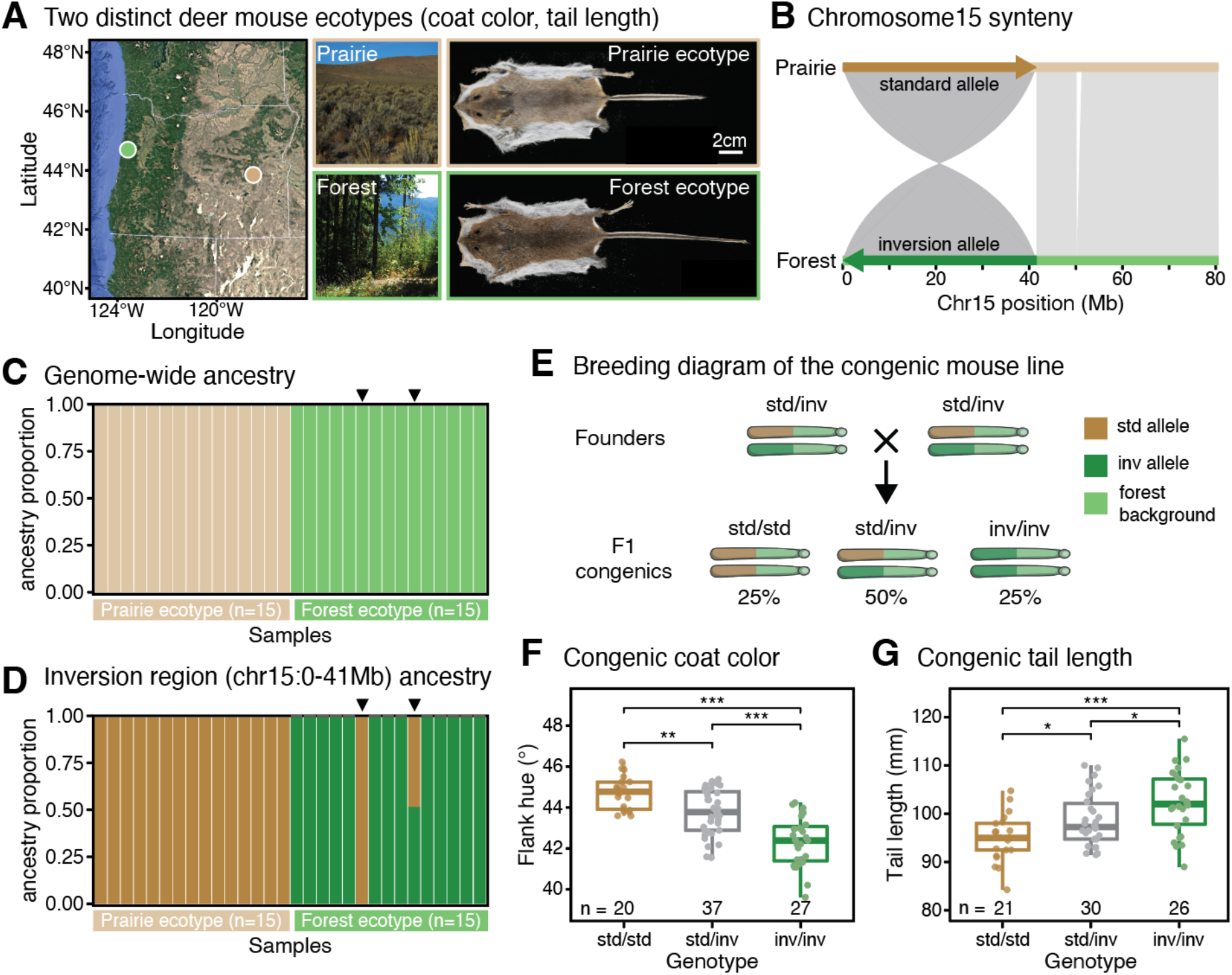
An inversion supergene is associated with multiple forest-adaptive traits. (**A**) Prairie and forest ecotypes of deer mice sampled from eastern and western Oregon, respectively. Photographs show typical habitats for each ecotype (photo credit: Emily Hager). Flat-skin images show a representative prairie mouse with a short tail and light coat, and a forest mouse with a long tail and dark coat. (**B**) Synteny plot of chromosome 15 between prairie and forest mouse genomes, showing the 41-Mb chromosomal inversion. (**C**) Genome-wide ancestry proportions of wild-caught prairie and forest mice used as founders of the laboratory colony. (**D**) Ancestry proportions across the inversion region (0-41Mb on chr15). Arrows indicate two forest individuals carrying the *standard* haplotype (one homozygous, one heterozygous). Ancestry data are from (*16*). (**E**) Breeding scheme for the congenic line. These mice share a common forest genomic background but differ at the inversion locus: *std* = standard haplotype, *inv* = inversion haplotype. Crosses between *std/inv* individuals produce sibling offspring of all three inversion genotypes: *std/std*, *std/inv* and *inv/inv*. (**F**) Effect of the inversion on coat color in congenic mice at P70-75, measured as flank hue (degrees); *P* < 0.05 (ANOVA). (**G**) Effect of the inversion on tail length in congenic mice at P70-75; *P* < 0.05 (ANOVA). Sample sizes are indicated for each genotype. Asterisks indicate \**P* < 0.05, \*\**P* < 0.01, \*\*\**P* < 0.001 (two-sided Welch’s t-tests). Boxplots show median (center line), interquartile range (box), and values within 1.5 × interquartile range (whiskers); points represent individual animals.

### Inversion drives phenotypic changes in a congenic background

Since forest and prairie mice are genetically differentiated (Fig. 1C), we isolated the effect of this inversion in a controlled genetic background. Although the inversion (*inv*) haplotype is highly prevalent in forest mice, it is not fixed (*16*). We therefore screened laboratory-reared offspring of wild-caught forest mice and identified two founders carrying the standard (*std*) haplotype in addition to *inv*/*inv* founders (Fig. 1D). Intercrosses generated sibling offspring representing all three inversion genotypes—*std/std*, *std/inv*, and *inv/inv*—within an otherwise homogeneous forest genetic background (Fig. 1E). Phenotypic analysis of these congenic animals confirmed strong effects of the inversion on both coat color and tail length. Homozygous *inv/inv* mice exhibited significantly darker coats than *std/std* animals, fully recapitulating the pigmentation differences between forest and prairie ecotypes (Fig. 1F, Fig. S1A). Similarly, *inv/inv* mice grew tails that were on average 7.3mm longer than those of s*td/std* animals, accounting for 21.8% of the tail length divergence between ecotypes (Fig. 1G, Fig. S1, B and C). Heterozygous *std/inv* animals had intermediate phenotypes for both traits, suggesting the effects of the inversion are additive (Fig. 1, F and G). Together, these results are consistent with previous estimates from an F_2_ intercross (*16*) and provide direct genetic evidence that the inversion functions as a supergene.

### Differential expression of Slc45a2 underlies the inversion’s effect on coat color

We first investigated how this inversion influences coat color. Like many mammals, deer mice transiently produce yellow pheomelanin instead of dark eumelanin during early anagen, generating a subapical agouti band along the hair shaft (Fig. 2, A and B) (*17, 18*). Variation in this banding pattern can markedly alter overall coat coloration (*19, 20*). Indeed, *inv/inv* mice possess a subtle agouti band with less pheomelanin deposition compared to *std/std* animals (Fig. 2B, Fig. S2A). By contrast, the overall proportion of banded hair, as well as the average number of mature melanocytes per hair follicle, were unaffected by the inversion (Fig. S2B-D). These results indicate that the inversion darkens coat color by specifically reducing the pheomelanin-to-eumelanin ratio during agouti band formation.

**Figure 2.**
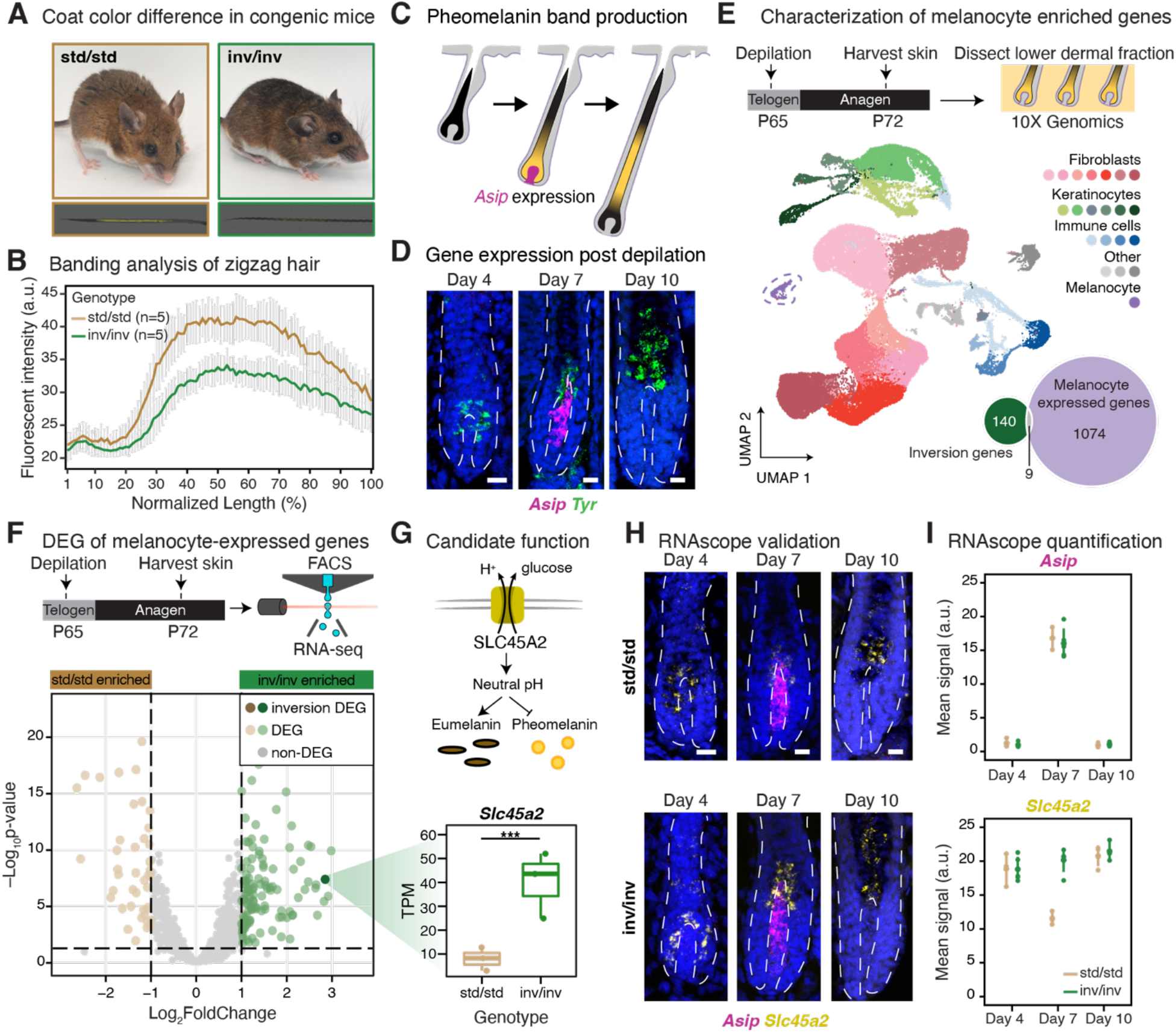
Identifying candidate genes underlying the inversion’s effect on coat color. (**A**) Representative images of adult *std/std* and *inv/inv* mice and the tip of their zigzag hairs, illustrating the agouti band. Autofluorescence (pseudo-colored in yellow) indicates pheomelanin deposition. (**B**) Quantification of autofluorescence intensities, normalized to the distance between the hair tip and the first zigzag bend. 5 mice per genotype were analyzed, with 10 hairs from each mouse. (**C**) Schematic of melanin production in the hair follicle. *Asip* expression in dermal papilla cells promotes switching from eumelanin (black pigment) to pheomelanin (yellow pigment). (**D**) RNAscope detection of *Asip* and melanocyte marker *Tyr* expression in hair follicles at 4, 7, and 10 days post depilation. Scale bars, 20 µm. (**E**) Experimental design for single-cell RNA sequencing: mice were depilated at P65 and anagen dorsal skin was collected at P72 during agouti band formation. UMAP visualization of skin cells from 3 *std/std* and 3 *inv/inv* samples, colored by cell type. Venn diagram depicts the intersection between inversion genes (n=149) and melanocyte-expressed genes (n=1,083). (**F**) Differentially expressed genes (DEGs) between FACS enriched *std/std* and *inv/inv* melanocyte samples. Dashed lines indicate thresholds of Log2FoldChange = ±1 and adjusted-*P* = 0.05. DEGs are highlighted in color; the DEG located within the inversion, *Slc45a2*, is highlighted in darker color, with its expression levels in 3 *std/std* and 3 *inv/inv* mouse melanocyte samples shown in the boxplot; TPM = transcripts per million. (**G**) Cartoon diagram illustrating the function of SLC45A2 in pigment production. (**H**) RNAscope validation of *Slc45a2* and *Asip* expression in the hair follicles of *std/std* and *inv/inv* mice at 4, 7, and 10 days post depilation. Scale bars, 20 µm. (**I**) Quantification of RNAscope signal of *Asip* and *Slc45a2* in the hair follicle. Mean fluorescent intensity is shown.

To identify candidates within the inversion that affect this banding phenotype, we first focused on genes carrying protein coding changes. The inversion encompasses 149 annotated genes, 29 of which harbor at least one nonsynonymous variant fixed between the *inv* and *std* haplotypes (Fig. S2E). Intersecting this list with genes known to affect pigmentation phenotypes in the laboratory mouse yielded a single candidate, *Slc45a2*, which carries an L75F substitution in the *inv* allele (Fig. S2F). However, multispecies sequence alignment revealed that both leucine and phenylalanine occur at position 75 across mammals despite strong conservation of the surrounding residues, implying this variant is unlikely to be disruptive (Fig. S2G). Moreover, an independent deer mouse population (*P.m.rufinus*) carrying the same inversion on chr15 lacks the L75F mutation but displays the same dark-coat phenotype, ruling out SLC45A2^L75F^ as the causal variant (Fig. S2H). Together, these observations point towards non-coding regulatory changes at the inversion driving the coat color effects.

To identify gene expression changes underlying pigmentation differences between *inv/inv* and *std/std* mice, we first established the timing of agouti band production during the hair cycle. Using RNAscope *in situ* hybridization, we found that *Agouti signaling protein* (*Asip*), which inhibits melanocortin 1 receptor (*Mc1r*) and promotes pheomelanin synthesis, is transiently expressed in the dermal papillae seven days post depilation (Fig. 2C and D) (*19, 21*). We therefore focused on this time point and performed single-cell RNA sequencing on three *std/std* and three *inv/inv* individuals to characterize gene expression patterns across anagen skin (Fig. 2E). We identified 21 cell populations, each represented at comparable proportions across genotypes, including two important cell-types involved in pigmentation: dermal papilla fibroblasts (*Nrg2*^+^/*Fgf10*^+^) and melanocytes (*Tyr*^+^/*Pmel*^+^) (Fig. 2E, Fig. S3, A-C). Dermal papilla fibroblasts showed minimal transcriptomic differences, with zero inversion genes being significantly differentially expressed between *std/std* and *inv/inv* mice (Fig. S4B). *Asip* expression was also unchanged in *inv/inv* mice, suggesting the pigmentation phenotype is mediated by genes acting downstream of the *Mc1r* pathway in melanocytes (Fig. S4, A and B). We therefore used the single-cell dataset to define a transcriptomic signature of pheomelanin-producing melanocytes and identified 1,083 genes that passed our expression threshold (see Methods), 9 of which are found within the inversion (Fig. 2E). However, since melanocytes represented only 0.9% of cells in our single-cell dataset, differential expression analysis for these cells was underpowered (Fig. 2E). As an alternative approach to enrich for melanocytes, we took advantage of the surface marker CD117 (c-Kit), which, at early anagen, labels melanocytes, melanocyte stem cells, mast cells and a subset of matrix keratinocytes (Fig. S4C). Using fluorescence-activated cell sorting (FACS) at this same time point for three *std/std* and three *inv/inv*, we performed RNA sequencing and focused on the melanocyte expressed genes. This revealed one inversion gene with significant *inv/inv* versus *std/std* differential expression: *Slc45a2* (Fig. 2F).

*Slc45a2* encodes a putative membrane-associated transporter that regulates melanosome pH and promotes eumelanin production (Fig. 2G) (*22, 23*). Consistent with this role, *Slc45a2* shows higher expression in melanocytes of *inv/inv* compared to *std/std* mice (Fig. 2F, H and I). Notably, RNAscope results indicated that this differential expression was transient: *Slc45a2* is expressed at comparable levels between *std/std* and *inv/inv* animals both before and after agouti band formation (Fig. 2H and I). In the *std* haplotype, *Slc45a2* expression appears to be coupled to *Mc1r* pathway activity, decreasing as expression of the *Mc1r* antagonist *Asip* increases during agouti band formation (Fig. 2I). In the *inv* haplotype, *Slc45a2* expression remains stable during the period of peak *Asip* expression, consistent with a regulatory mutation that disrupts this transcriptional coupling (Fig. 2I). This maintained *Slc45a2* expression likely creates a condition less favorable for pheomelanin production, thereby increasing the relative abundance of eumelanin and ultimately darkening the coat (Fig. S4D).

### The inversion prolongs postnatal tail growth through downregulation of Npr3

We next examined how the inversion influences tail length. The mouse tail comprises a series of articulated caudal vertebrae (CVs) that progressively transition from stout, spinous elements proximal to the sacrum into slender, hourglass-shaped distal segments (Fig. 3A) (*24*). A long tail could therefore arise through several mechanisms: an increase in CV number, an increase in vertebral length, or a homeotic transformation of vertebral identity (*25–27*). X-ray analysis of 10-week-old congenic mice showed that, relative to *std/std* individuals, *inv/inv* mice develop longer CVs throughout the tail while maintaining the same CV count (Fig. 3, A-D, Fig. S5A). The effect is most pronounced across the middle section of the tail, spanning CV_4_ to CV_24_ (Fig. 3B). Notably, the characteristic “crescendo–decrescendo” pattern of CV proportions along the anterior–posterior axis remains unchanged, indicating preserved positional identities (Fig. 3B) (*28, 29*). This bone elongation phenotype was also tail-specific: thoracic, lumbar and sacral vertebrae, as well as long bones such as the tibia, showed no differences in length between *inv/inv* and *std/std* mice (Fig. S5, B-E, Fig. S8B).

**Figure 3.**
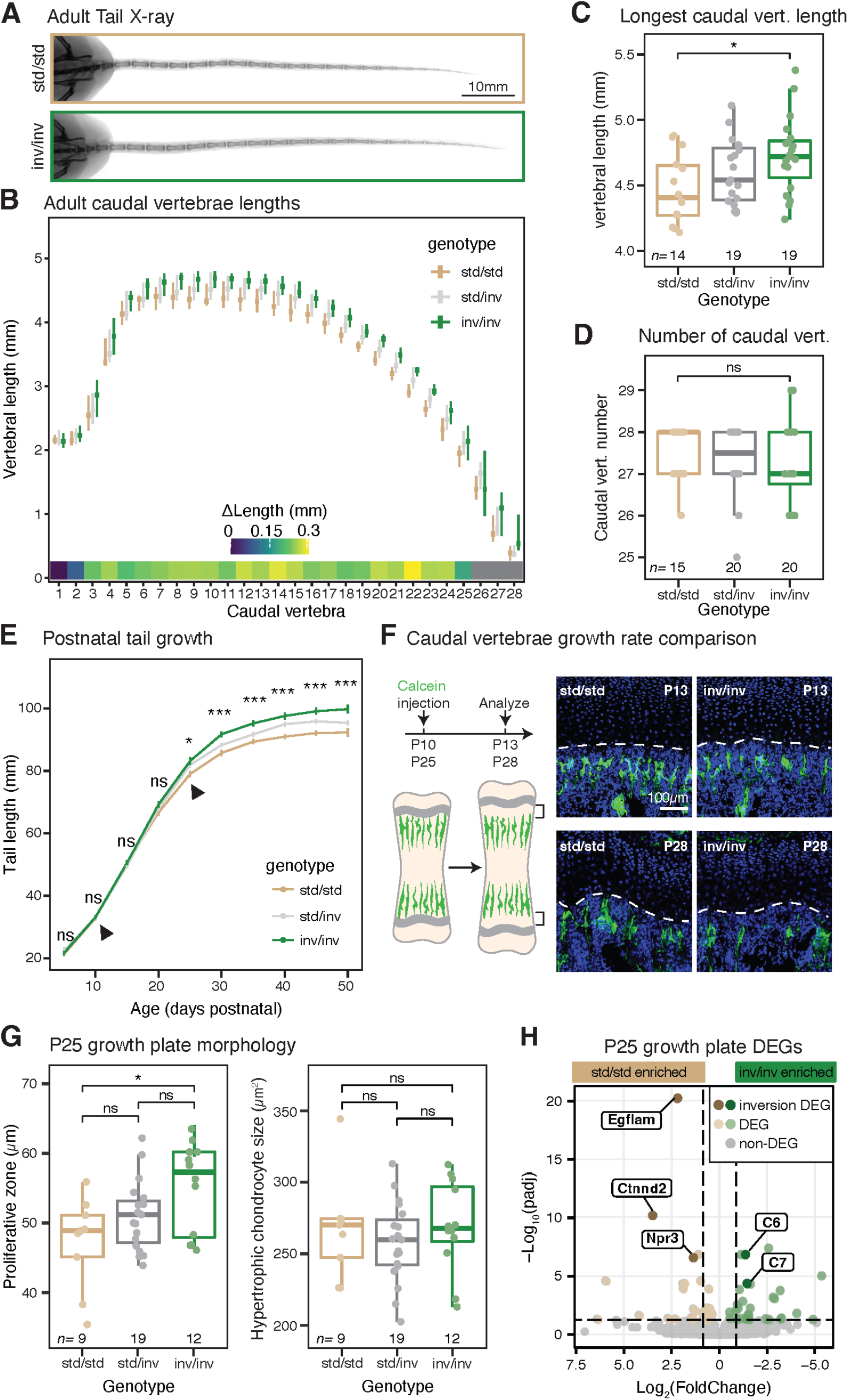
Identifying candidate genes underlying the inversion’s effect on tail length. (**A**) Representative X-ray images of tails from adult *std/std* and *inv/inv* mice. (**B–D**) Quantification of tail skeletal morphology from X-ray images of congenic mice at P70–75. (**B**) Length of individual caudal vertebrae along the tail. A linear model (vertebral length ∼ genotype + caudal vertebra number) detected a significant effect of genotype (*P* < 0.001). Heatmap below shows differences in vertebral length between genotypes. (**C**) Length of the longest caudal vertebra. (**D**) Total number of caudal vertebrae. Sample sizes for each genotype are indicated. Asterisks: \**P* < 0.05 (two-sided Welch’s *t*-tests). (**E**) Tail growth trajectories in congenic mice (5-day average). Sample sizes: *n* = 17 (*std/std*), 32 (*std/inv*), and 23 (*inv/inv*). Significance is shown for each time point. Asterisks: \**P* < 0.05, \*\*\**P* < 0.001 (ANOVA). Arrows indicate time points selected for follow-up analyses. (**F**) Schematics of the calcein pulse-chase experiment and representative images of CV_6_ caudal vertebral growth plates showing calcein incorporation (green) at P13 and P28, 3 days post injection. (**G**) Quantification of CV_6_ growth plate features at P25. Left, thickness of the proliferative chondrocyte zone, *P* < 0.05 (ANOVA). Right, size of hypertrophic chondrocytes, *P* > 0.05 (ANOVA). Symbols: ns*=P* > 0.01; \**P* < 0.01 (two-sided Welch’s *t*-tests). (**H**) RNA sequencing of caudal vertebral growth plates at P25 from 6 *std/std* and 5 *inv/inv* mice. Differentially expressed genes (DEGs) are highlighted in color; the 5 DEGs located within the inversion are shown in darker colors and labeled. Dashed lines indicate thresholds of Log2FoldChange = 1 and adjusted-*P* = 0.05.

We then asked how this difference in CV length arises during development. Whole-mount skeletal staining of P0 pups suggested that the tail length disparity emerges postnatally (Fig. S6, A and B). Indeed, *std/std* and *inv/inv* mice exhibited indistinguishable tail growth trajectories during the first three weeks after birth, but diverged around P25, when growth slowed in *std/std* mice and ceased by six weeks of age––approximately two weeks earlier than in *inv/inv* animals (Fig. 3E). Consistent with these findings, a pulse–chase assay using calcein, a fluorescent dye that incorporates into skeletal tissue as it is mineralized, confirmed a clear difference in vertebral growth rates at P25 but not at P10 (Fig. 3F, Fig. S6, C and D). Because the rate of longitudinal extension of CVs depends on the degree of cell swelling of hypertrophic chondrocytes as well as the level of proliferative activity in the growth plates, we examined CV growth plate morphology at P25 (Fig. S7, A and B). Histological analysis revealed increased heights of both proliferative and hypertrophic zones in *inv/inv* CV_6_ growth plates, with no change in hypertrophic chondrocyte size (Fig. 3G, Fig. S7, C-E). Concordantly, *inv/inv* growth plates contained more *Ki67*⁺ proliferative chondrocytes, *p57*⁺ hypertrophic chondrocytes, and *Sox9*⁺; *Ki67*⁻ resting chondrocytes, whereas the density of hypertrophic chondrocytes remained unchanged (Fig. S7, F-H). Together, these findings demonstrate that the inversion prolongs postnatal tail growth by sustaining proliferative activity within caudal vertebral growth plates.

To pinpoint candidate gene(s) underlying the inversion’s effect on CV length, we adopted a two-pronged strategy combining transcriptomic profiling with forward genetic mapping. RNA sequencing of dissected congenic CV growth plates at P25 identified 5 differentially expressed genes within the inversion (Fig. 3H). Because the inversion does not affect appendicular growth, tibia growth plates were profiled in parallel as an internal negative control, allowing us to prioritize genes with tail-specific expression differences (Fig. S8, A-C). Intersecting these datasets narrowed the list to only three genes—*Egflam*, *Ctnnd2*, and *Npr3*—that showed CV-specific differential expression between *inv/inv* and *std/std* mice.

As a complementary approach, we fine-mapped tail length within the inversion by taking advantage of a deer mouse population (*P.m.rufinus*) from New Mexico carrying a unique *inv* haplotype (Fig. 4A). Through long-read sequencing based *de novo* genome assembly, we confirmed that the same inversion on chr15 segregates in this population at 50% allele frequency (Fig. 4, B and C). Unexpectedly, *std/std* and *inv/inv P.m.rufinus* mice differed in coat color but not in tail length, suggesting that the mutation(s) responsible for tail elongation are absent in the *rufinus inv* haplotype (Fig. 4D). We hereafter refer to the *rufinus* haplotype as *inv^t^* and the congenic haplotype as *inv^T^* to reflect their phenotypic differences. This fortuitous finding enabled a forward genetic cross between *inv^T^/inv^T^* (forest) and *inv^t^*/*inv^t^*(*rufinus*) mice, generating a mapping population of 373 F_2_ hybrids that exhibited a range of intermediate tail lengths and recombination within the inversion region (Fig. 4, E and F). QTL mapping identified three loci with significant effects on tail length, together explaining 22.5% of the phenotypic variance (Fig. 4G, Fig. S9). The strongest effect QTL localized to a 5.7-Mb region within the chr15 inversion, with each *inv^T^* allele increasing tail length by 2.5mm on average (Fig. 4, H and I). Out of the 33 genes within this fine-mapped tail length region, only *Npr3* was differentially expressed in CV growth plates between congenic mice (Fig. 4, J-K).

**Figure 4.**
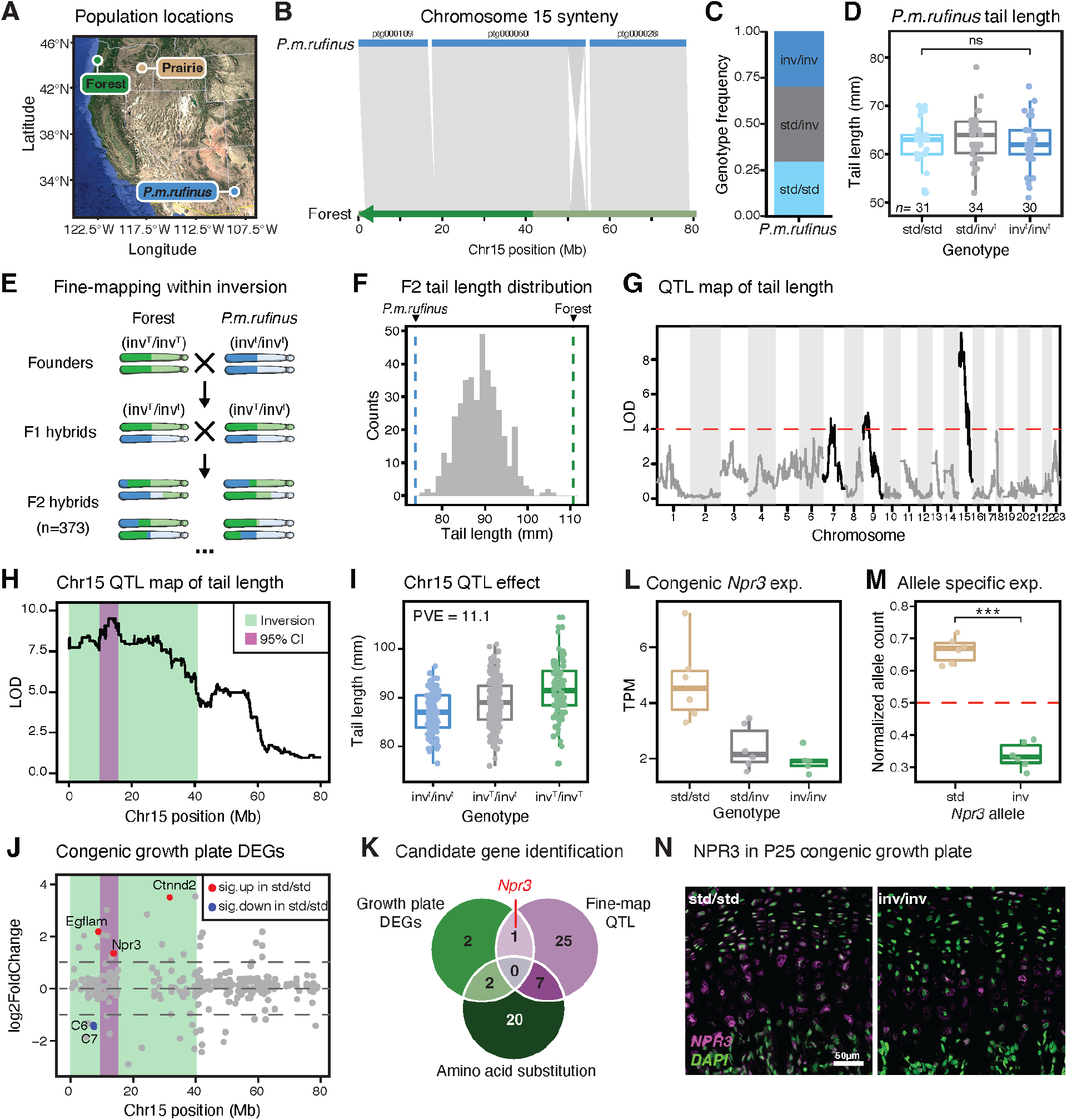
Fine-mapping tail length within the inversion. (**A**) Location of the *P.m.rufinus* population in New Mexico, with focal forest and prairie populations also indicated. (**B**) Synteny plot between the forest mouse chromosome 15 and the *P.m.rufinus* contig-level long-read genome assembly. Contig ptg000060l spans the inversion breakpoint, indicating that *P.m.rufinus* carries the same inversion. (**C**) Frequency of the inversion genotypes in *P.m.rufinus*. (**D**) Tail length measurements for *P.m.rufinus* mice with sample sizes indicated. No significant difference in tail length was detected between genotypes (two-sided Welch’s t-test). (**E**) Schematic of the forward genetic cross between forest *inv^T^/inv^T^* mice and *P.m.rufinus inv^t^/inv^t^* mice. *inv^t^* denotes the inversion haplotype without effects on tail length, whereas *inv^T^* denotes the inversion haplotype associated with tail length effects. Chromosome colors indicate local ancestry, with the inversion region shown in darker shades. (**F**) Histogram of F2 tail length. Mean tail lengths of *inv^t^/inv^t^* New Mexico mice and *inv^T^/inv^T^* forest mice are indicated by blue and green lines, respectively. (**G**) QTL map of tail length in F2 hybrids (*n* = 373). Genome-wide significance threshold from 1,000 permutations is indicated by the red dashed line. (**H**) A zoomed-in view of the QTL peak on chromosome 15. The inversion region is highlighted in green, and the 1-LOD confidence interval for the fine-mapping QTL is highlighted in purple. (**I**) Effect of the chromosome 15 QTL in F2 hybrids, grouped by genotype at the peak marker (*inv^t^*= *P.m.rufinus* allele; *inv^T^* = forest allele). PVE, percent variance explained. (**J**) Genomic positions of differentially expressed genes (DEGs) between *std/std* and *inv^T^/inv^T^* CV_6_ growth plates along chromosome 15. Green and purple shaded regions indicate the inversion and fine-mapping QTL intervals, respectively. Significant DEGs are shown in blue (lower expression in *std/std* mice) and red (higher expression in *std/std* mice). (**K**) Identification of candidate tail-length genes within the inversion: among the 5 growth plate DEGs, 1 gene (*Npr3*) falls within the chromosome 15 fine-mapped QTL interval. (**L**) *Npr3* expression in CV_6_ growth plates at P25; TPM = transcripts per million. Sample sizes: n = 6 (*std/std*), 6 (*std/inv*), and 5 (*inv/inv*). (**M**) Allele-specific expression of *Npr3* measured in CV_6_ growth plates of 6 *std/inv* mice. Normalized allele counts, averaged across 12 allele-informative SNPs, are shown for *std* and *inv Npr3* alleles. Symbols: \*\*\**P* < 0.001 (paired t-test). (**N**) Representative immunostaining of NPR3 (magenta) in chondrocytes of P25 vertebral growth plates, with DAPI shown in green.

*Npr3* encodes natriuretic peptide receptor C, a decoy receptor for C-type natriuretic peptide (CNP), a key signaling molecule in endochondral ossification (*30, 31*). In both laboratory mouse and human patients, functional disruption of *Npr3* enhances CNP signaling, expands the proliferative and hypertrophic zones of the growth plate, and produces a dramatic skeletal overgrowth phenotype (*32–35*). Consistent with this mechanism, congenic *inv/inv* mice exhibited significantly reduced *Npr3* expression in CV growth plates relative to *std/std* mice, a result independently validated by RNAscope and immunostaining (Fig. 4, L and N, Fig. S10, A-B). In addition, allele-specific expression analyses in *std/inv* heterozygotes revealed that *Npr3* expression is strongly biased toward the *std* haplotype (Fig. 4M, Fig. S10C). By contrast, *Npr3* expression was unaffected by genotype in tibia growth plates (internal control), suggesting that the underlying mutation resides in tail-specific *cis* regulatory elements (Fig. S8D). Together, these findings indicate that reduced *Npr3* expression is likely responsible for the inversion’s effect on tail length, with elevated CNP signaling sustaining chondrocyte proliferation and extending the postnatal growth period of the tail (Fig. S10D).

### Evolution of the inversion supergene

After defining how the inversion influences two locally adaptive forest traits and identifying candidate genes underlying these effects, a key remaining question was when the mutations affecting coat color and tail length arose relative to the inversion itself. To address this, we reconstructed the supergene’s evolutionary history by first genotyping for the inversion across the range of deer mice. Using whole-genome resequencing data from 14 wild populations, we inferred inversion genotype in 692 individuals and found that this inversion is widespread across western North America, segregating in 7 populations and reaching its highest frequencies along the forested northwestern coast (Fig. 5A). We then assessed the inversion’s phenotypic effects in four focal populations and found that, while the coat-color effect is broadly conserved, the tail-length effect is geographically restricted, indicating that the inversion functions as a supergene only in part of its range (Fig. 5B).

**Figure 5.**
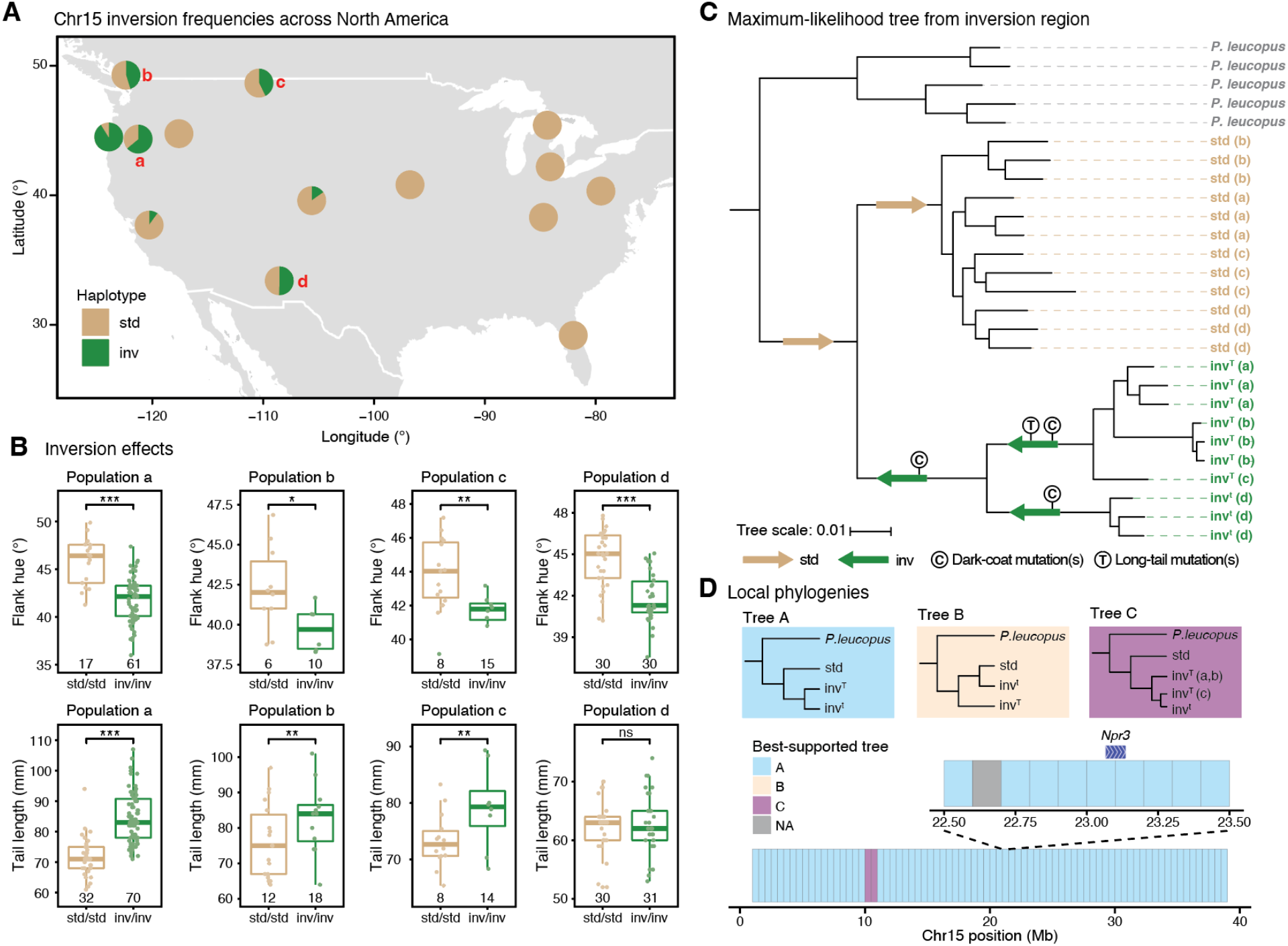
Evolutionary history of the inversion supergene. (**A**) Map of inversion frequencies across the species range. Pie charts show inversion frequencies, with *inv* in green and *std* in tan. Red letters denote populations analyzed in (**B**). (**B**) Phenotypic effects of the inversion in four populations in which the inversion is polymorphic. Coat color effects, measured as flank hue (degrees) are shown above, and tail length effects are shown below. Symbols: ns = *P* > 0.05; \**P* < 0.05; \*\**P* < 0.01; \*\*\**P* < 0.001 (two-sided Welch’s t-tests for coat color; linear models for tail length, see Methods). Boxplots indicate the median (center line), the 25th and 75th percentiles (box limits), whiskers extend to the largest or smallest value within 1.5 times the interquartile range, and points represent individual samples, with sample sizes indicated below. (**C**) Maximum-likelihood tree inferred from SNPs in the inversion region (chromosome 15: 0–41 Mb). *P. leucopus* was used as the outgroup. The population of origin for each haplotype is shown in parentheses. *inv^T^*and *inv^t^* denote the inversion haplotypes with or without tail length effects, respectively. Arrows on the tree show the most likely haplotype in the shared ancestor. (**D**) Local gene trees across the inversion region inferred from 500-kb non-overlapping windows. Inset shows local gene trees for 100-kb non-overlapping windows spanning the *Npr3* region.

We next constructed a maximum-likelihood tree based on SNPs within the inversion to resolve relationships among haplotypes (Fig. 5C). The *std* haplotype represents the ancestral state and is also found in the outgroup species *Peromyscus leucopus*. All inversion haplotypes from populations a–d formed a monophyletic clade, indicating descent from a common ancestral inversion haplotype associated with dark coat-color (Fig. 5C). Within this clade, *invᵀ* haplotypes (with tail-length effects) diverged from *invᵗ* haplotypes (without tail-length effects), consistent with either loss of ancestrally acquired tail-length mutation(s) in the *invᵗ* lineage or gain of *de novo* tail-length mutation(s) in the *invᵀ* lineage––the latter being the more parsimonious explanation (Fig. 5C). A remaining possibility, however, is that *invᵗ* lost tail-length mutation(s) through a rare double recombination event with a *std* haplotype. To test this, we reconstructed gene trees in 500-kb windows across the inversion. These trees consistently supported shared ancestry between *invᵀ* and *invᵗ*, with no evidence that *invᵗ* is more closely related to *std* haplotypes in any window (Fig. 5D). Because the inversion most likely affects tail length through *cis*-regulatory changes of *Npr3*, we further scanned the surrounding region of *Npr3* at a 100-kb resolution and again found only support for shared ancestry between *invᵀ* and *invᵗ* (Fig. 5D). These results argue against recombination-mediated loss of the *Npr3* allele in *inv^t^*; furthermore, they suggest that *de novo* mutation within the inversion, rather than standing variation from populations carrying the *std* haplotype, is likely responsible for the tail length effects. Together, our findings support a model in which the inversion first arose and captured—or rapidly accumulated—mutation(s) affecting coat color, followed by the origin of tail-length mutation(s) in a subset of inversion haplotypes.

## Discussion

The molecular dissection of the chr15 inversion in deer mice revealed novel mechanisms of morphological adaptation and the evolutionary route by which a supergene can emerge. As a core component of the melanosome, SLC45A2 has been implicated in natural color variation across diverse mammalian species, including horses, tigers, gorillas, and humans (*36–40*). In most documented cases, however, these pigmentation differences are associated with coding variants that constitutively alter SLC45A2 protein function and consequently produce coordinated changes in hair, skin, and eye pigmentation. Here, through systematic phenotypic and transcriptomic analyses, we found that the effect of the chromosome 15 inversion on deer mouse coat color is not caused by a coding change in SLC45A2, but rather by a transient expression difference in early anagen. Regulatory mutations of this type are harder to detect yet may be favored over coding changes because they can fine-tune melanogenesis in a temporally and tissue-specific manner, enabling adaptive modification of pelage color while minimizing pleiotropic effects on pigment production in other tissues.

Unlike coat color, skeletal evolution is typically thought to involve a more complex polygenic architecture (*41*). Unexpectedly, our fine-mapping and transcriptomic analyses implicate tail-specific downregulation of a single gene, *Npr3*, as the primary driver of caudal vertebral elongation in *inv/inv* mice. Echoing our finding, a recent report linked differential expression of *Npr3* to the extreme disproportionate growth of tail vertebrae in jerboa, suggesting this locus may be repeatedly targeted during evolution (*29*). Notably, *Npr3* has also been associated with increased tibia length in the artificially selected *Longshanks* mouse, where tail length remains unchanged (*42*). These results suggest that multiple distinct regulatory elements control *Npr3* expression in skeletal tissue of distinct developmental origin. *Npr3* therefore represents an illustrative example of regulatory modularity in skeletal evolution. Key next steps will be to identify such regulatory elements and causal mutations altering *Slc45a2* and *Npr3* expression.

A longstanding question in evolutionary biology is how chromosomal inversions establish and persist as polymorphisms (*1, 4*). Inversions are generally expected to be disfavored because they can disrupt gene structure and regulatory architecture at their breakpoints, impair meiosis in heterozygotes, and accumulate deleterious mutations as reduced recombination limits purifying selection. Theory therefore predicts that stably maintained inversions should be those that capture—or generate—multiple adaptive loci at their origin, such that their cumulative selective advantage as a supergene outweighs these costs (*5, 6, 12*). Our results support an alternative model: a large inversion that established first and acquired supergene properties later through sequential accumulation of adaptive variants. The *Peromyscus* chr15 inversion was likely maintained initially at intermediate frequency due to selection on its coat color effect, given the substantial fitness benefit associated with pigmentation in natural populations (*43, 44*). The inversion’s terminal position on chr15 may also reduce inversion-loop formation in *std/inv* heterozygotes, thereby weakening underdominant effects and allowing its persistence (*45*). The subsequent emergence of tail-length mutation(s) near *Npr3* further increased the inversion’s selective advantage by coupling two locally adaptive alleles: *Slc45a2* and *Npr3* are 0.8 Mb apart, a distance at which we predict strong fitness gain from the inversion’s suppression of recombination in the face of gene flow (*16*). This additional fitness benefit likely then drove the inversion to near fixation in the northwestern coastal forests. In summary, our findings suggest that inversions need not originate with a favorable combination of adaptive alleles but can instead provide a genomic substrate on which beneficial mutations accumulate and are subsequently maintained together as a supergene.

## Author contributions

O.S.H., S.H. and H.E.H. conceived of the study. S.H. and O.S.H. performed the experiments and analyses, and wrote the manuscript together with H.E.H.

## Acknowledgements

The authors thank Ya-Chieh Hsu, Sarah Kocher, Janet Song, and Cliff Tabin for providing helpful feedback on the manuscript. The authors thank Chris Kirby, Sade McFadden and Isobel Smith for assistance with animal husbandry and data collection. The Bauer Core Facility at Harvard University provided library preparation and sequencing services. Computational analyses were run on the Odyssey and Cannon clusters supported by the Faculty of Arts and Sciences Research Computing Group at Harvard University. The authors thank the Museum of Southwestern Biology (University of New Mexico) and Museum of Comparative Zoology (Harvard University) for providing specimens used in this study.

## Data, code, and materials availability

Associated data and code will be uploaded to Dryad, NCBI and GitHub prior to publication.

## AI statement

AI was not used in this study for analyses or writing.

## Supplementary Materials

Materials and Methods

Figs. S1 to S11

Tables S1 to S2

## Supplementary Materials

## Materials and Methods

### Ethics statement

Experiments were approved by the Harvard University Institutional Animal Care and Use Committee under the protocols 11-05 and 27-16 and were conducted in accordance with National Institutes of Health regulations governing the humane treatment of vertebrate animals.

### Generation of the congenic mouse line

Three custom Taqman SNP genotyping assays (Life Technologies) spanning the 41-Mb inverted genomic region were designed to distinguish between the *inv* and *std* alleles at locations 14,131,681, 23,939,629 and 29,639,255 on chromosome 15 (Table S1). These SNPs were selected as diagnostic SNPs differentiating the inversion and standard haplotypes based on whole-genome sequencing data of mice (n=30) from the forest and prairie ecotypes (*16*). Genomic DNA was extracted from ear tissue using the Maxwell RSC DNA kit (Promega) following manufacturer’s instructions. All Taqman genotyping reactions were performed and analyzed using a Mastercycler RealPlex^2^ system (Eppendorf), with the following PCR parameters: 95 °C for 10 minutes followed by 40 cycles of 95 °C for 15 s, 60 °C for 1 minute.

Since the chromosome 15 inversion is not fixed in the forest population, two forest (*P.m.rubidus*) mice harboring the *std* haplotype (1 *std/inv* forest mouse and 1 *std/std* forest mouse) served as founders of the forest mouse lab colony (Fig. 1D), resulting in both *inv* and *std* haplotypes segregating within the forest lab colony. Using the taqman SNP genotyping assays described above, we identified 6 male and 6 female heterozygous mice (*std/inv*) in our established lab forest mouse colony and paired these mice as founders for a congenic mouse line. The genotypes of subsequent generations were determined using the same three diagnostic Taqman SNP assays.

### Coat color measurement

Coat color was measured using a FLAME UV-VIS spectrometer with a pulsed xenon light source, a 400µm reflectance probe, and OceanView software (Ocean Optics). We obtained the reflectance spectra at 5 spots along the dorsal midline as well as the flank region. A custom R script was then used to calculate the brightness, hue, and saturation values in the range of 400-700 nm with a 1-nm bin width. For each trait, we calculated the median values from the reflectance spectra measurements. For the congenic mice, we measured coat color at post-natal day 70 (P70), when the adult coat is established.

### Hair banding analysis

We prepared flat skins and then used 1-mm hair punches to collect hairs from the dorsal region of the mouse skins. Consistent locations of the dorsal regions were used for hair collection through ruler measurements of the flat skins. We then selected 10 zigzag hairs (the major pigmented hair type found within mouse coats) per individual from hair plucks of 5 *std/std* and 5 *inv/inv* mice and imaged pheomelanin-based autofluorescence using a Zeiss Axio imager microscope following a published protocol (*20*). We then used Fiji (ImageJ) to quantify the linear autofluorescence intensity along the first distal zigzag bend and normalized the distances from hair tip to the first bend to account for hair length differences.

### SLC45A2 amino acid conservation

Multi-species SLC45A2 amino acid sequences were downloaded from the NCBI Protein Database and aligned using Clustal Omega (https://www.ebi.ac.uk/jdispatcher/msa/clustalo). Alignment was then plotted using the ggmsa package in R (*46*).

### Skin single-cell RNA sequencing

Adult congenic mice of both genotypes (3 *std/std* and 3 *inv/inv*) were depilated at telogen using cold wax strip (Veet) to induce a round of synchronized hair growth. Back skins were harvested 8 days post depilation. Skin associated white adipose tissue and panniculus carnosus muscle were carefully stripped off using a pair of fine tweezers. The samples were then placed dermal side down and digested using 1mg/mL Collagenase I (Sigma-Aldrich, C2674) diluted in Hanks’ Balanced Salt Solution (ThermoFisher, 24020117) for 30 mins at 37°C. A clean, sharp razor blade was used to scrape off the lower dermal fraction containing the hair bulbs and dermal papillae into fresh Collagenase I solution and digested for another 30 mins at 37°C with gentle shaking. Next, the samples were centrifuged at 300g, 4°C for 10 mins to collect the cell pellet and further digested in 0.25% Trypsin-EDTA (ThermoFisher, 25200056) for 30 mins at 37°C with gentle shaking. The samples were then passed through a 100µm Falcon® Cell Strainer (Sigma-Aldrich, CLS352360) and centrifuged at 300g, 4°C for 10 mins. The cell pellets were resuspended in 1x calcium and magnesium free PBS (ThermoFisher, 70011044) supplemented with 5% Fetal Bovine Serum (ThermoFisher, A5256701). Finally, the samples were pipetted through 40µm Flowmi® Cell Strainers (Sigma-Aldrich, BAH136800040) to obtain single cell suspensions. Single-cell RNA-sequencing libraries were generated from approximately 15K cells per sample with the 10X Genomics Chromium Next GEM 3’ Kit (v3.1) per manufacturer’s protocol. Pooled libraries were sequenced on a NovaSeq X 25B flow cell (Illumina).

Single-cell RNA-sequencing data were analyzed using Seurat v4 (*47*). Cells were first filtered based on nCount_RNA value between 500 and 10,000 using the *subset* function. All samples were subsequently downsampled to a maximum of 11,000 cells/sample to balance the number of cells per condition. Clustering was performed using PCA for dimensionality reduction followed by *FindNeighbors* with 20 dimensions and reduction=integrated.dr and *FindClusters* with resolution=0.3. The resulting 21 clusters were plotted with *RunUMAP* with reduction=integrated.dr. Feature plots were generated using the *scCustomize* package. To identify differentially expressed genes in the clusters of interest between *inv/inv* and *std/std* mice, we performed a pseudobulk analysis to aggregate gene expression by sample. We used *AggregateExpression* to pseudobulk gene expression by sample and then performed differential expression analysis with *DESeq2* using *FindMarkers* for each cell cluster of interest. We identified significant DEGs in the *Asip+* cluster using an adjusted p-value threshold of 0.05 and a log2FoldChange threshold of 1. To identify genes consistently expressed in mature melanocytes, we required the gene to have at least 1 count in more than 25% of transcriptionally defined melanocytes.

### Melanocyte fluorescence-activated cell sorting (FACS) and RNA sequencing

To enrich for melanocytes, we harvested the dermal fraction from the back skin of 3 *inv/inv* and 3 *std/std* animals 8 days post depilation and prepared single cell suspension as described above. The samples were then incubated with a CD117/c-kit-APC-Cy7 antibody (BioLegend 135136, 1:400) for 30 mins on ice. FACS experiments were performed on a BD FACSAria^TM^ II+ sorter. After sorting, cells were lysed directly into Trizol LS Reagent (Invitrogen). RNA was isolated using the RNeasy Micro Kit (Qiagen) and the QIAcube according to the manufacturer’s instructions. RNA was quantified using the TapeStation (Agilent) and then RNA libraries were prepared using the Kapa mRNA Hyperprep kit. Quality control on RNA libraries was performed with the TapeStation (Agilent) and qPCR. RNA libraries were sequenced as 2x150bp paired-end reads on a NovaSeq X platform.

To identify melanocyte differentially expressed genes, we aligned the sequencing reads to the *P. maniculatus* transcriptome using STAR (*48*). We created a STAR index using the NCBI gene annotation for *P. maniculatus*; we then aligned the RNAseq fastq files to the transcriptome, using quantMode of TranscriptomeSAM and outputting unsorted BAM files. We quantified transcript abundances using RSEM (*49*); we created an RSEM index and then ran rsem-calculate-expression with default parameters. Finally, we identified *inv/inv* versus *std/std* differentially expressed genes using DESeq2 (*50*). We restricted the differential-expression analysis to genes expressed in melanocytes from the single-cell dataset. We used a significance threshold of 0.05 for the adjusted p-value and a log2FoldChange threshold of 1 to identify differentially expressed genes.

### Tail length measurements

For each congenic mouse, we performed standard morphological measurements immediately following euthanasia at P70. We measured total length of the mouse (nose to tail tip), tail length and weight. To investigate vertebral morphology, we used a digital X-ray system (Varian Medical Systems, Inc.) in the Harvard Museum of Comparative Zoology Digital Imaging Facility to obtain X-ray images of the congenic mouse specimens. We imaged the whole specimen, including the tail, for each mouse. We mounted specimens such that the anterior-posterior and medio-lateral axes were parallel to the imaging plane. We then measured the caudal, sacral and rostral vertebrae with Fiji/ImageJ, and included a standard for scale. We also measured tail growth over time in the congenic mice. Starting at post-natal day 5, we measured tail length in live congenic mice every 2 days until post-natal day 50, and averaged tail length by sample over 5-day windows. Tibia lengths were also measured in congenic mice at post-natal day 1 (P1), day 25 (P25) and in adult mice.

### Skeletal staining

Skeletal staining (Alcian blue/Alizarin red) was carried out on P1 carcasses following a published protocol (*51*) with the following modifications: fixed carcass was stained in 0.005% Alcian blue at 4 °C for 6 hours to reduce the background; after staining, the samples were cleared in 50% glycerol+0.5% KOH for up to a week before transferring into 100% glycerol for long term storage and imaging. The samples were imaged under a Zeiss Discovery v8 stereoscope with an Axiocam 305 color camera and analyzed using ZEN software.

### Tail vertebrae growth speed measurement

Calcein (Sigma-Aldrich, C0875-5G) was injected intraperitoneally at 15mg/kg into pups of the age P10 and P25, respectively. Tail tissue was harvested 3 days post injection and fixed in 4% PFA at 4°C overnight. Cryosectioning was performed using a Leica CM3050 S cryostat with a thickness of 30µm. The sections were then air dried and counterstained using DAPI and imaged under a Leica LSM880 confocal microscope. The distance from the Calcein front to the edge of the chondro-osseous junction was measured and divided by 2 to calculate daily growth speed.

### Histology, immunostaining and RNAscope

To prepare skeletal samples for histological analysis, the sixth caudal vertebra (C6) was dissected from post-natal day 25 (P25) animals and fixed overnight at 4°C with 4% paraformaldehyde in 1xPBS. The bone samples were then decalcified in 10% EDTA for 1 week at room temperature and embedded in paraffin. Transverse sections through the center of the vertebrae were collected at 5 µm thickness and mounted on Superfrost plus microscope slides (Fisher Scientific). Safranin-O/Fast Green staining was carried out following a standard protocol.

To prepare skin samples for immunostaining, we harvested dorsal skin from mice at desired hair cycle stage. The samples were flattened on filter papers dermal side down and fixed in 4% paraformaldehyde in 1xPBS for 1 hour at room temperature with gentle agitation. Fixed samples were washed 3 times in 1x PBST, 20 mins each, and incubated overnight in 30% sucrose solution before embedding in OCT solution for cryosection. Cryosection was performed using a Leica CM3050 S cryostat at a thickness of 30µm and mounted on Superfrost plus microscope slides (Fisher Scientific).

For immunostaining, bone sections were dewaxed and rehydrated into 1x PBS solution. Antigen retrieval was carried out in 10mM Sodium Citrate solution with 0.05% Tween-20 at 65°C overnight. Samples were then incubated in primary antibodies diluted in blocking buffer (5% Normal Goat Serum, 1% BSA in 1xPBS) at 4°C overnight. Skin sections were washed 3 times in 1X PBST and incubated with 0.3% H_2_O_2_ in methanol at -20°C for 30 mins before proceeding with the standard immunostaining protocol. The following primary antibodies were used: Rat α-Ki-67 (1:200, Thermo Fisher, 14-5698-82), Rabbit α-Sox9 (1:200, Abcam, ab185966), Mouse α-Pax3 (1:200, DSHB), Rabbit α-PMEL (1:400, Abcam, ab137078), Mouse α-Npr2 (1:200, Thermo Fisher, H00004882-M02), Mouse α-Npr3 (1:100, Thermo Fisher, MA5-25065) and Mouse α-p57 (1:200, Santa Cruz, sc-56341). The following secondary antibodies were used: Goat anti-Rabbit IgG Alexa-Fluor 555 (1:1000, Thermo Fisher, A-21428), Goat anti-Mouse IgG Alexa-Fluor 555 (1:1000, Thermo Fisher, A-21422) and Goat anti-Rat IgG Alexa-Fluor 555 (1:1000, Thermo Fisher, A-21434). All slides were counter stained with DAPI (1:10000) as a reference signal. Images were acquired using either Zeiss Axio scan Z1 slide scanner or Zeiss LSM880 confocal microscope. Image analysis was performed using Fiji (ImageJ, v2.5.0). For vertebral length measurement, the average distance between the anterior and posterior epiphysis was measured at the center of each bone section. Growth plate thickness was measured based on Safranin-O signal, with at least 10 measurements taken across the center to account for tissue curvature. The size of hypertrophic chondrocytes was measured using the particle analysis tool with a manually defined measurement window. Distinct molecular markers (Ki67 for proliferative chondrocytes and p57 for hypertrophic chondrocytes) were utilized to visualize the position and density of different cell types on consecutive tissue sections. The average distance between the anterior and posterior edge of intensity-adjusted staining signal was calculated. Linear density of each cell type was calculated as the number of cells averaged by the curvature length of the growth plate within the measurement window.

RNAscope was carried out using the RNAscope multiplex fluorescent detection kit V2 (ACD Bio.) following manufacturer’s instructions. The following RNAscope probes were used: Pman-Agouti-C1 (1305261-C1), Pman-Tyr-C2 (1305281-C2), Pman-Slc45a2-C2 (1334061-C2), Pman-Npr3-C2 (1578651-C2) and Pman-Npr2-C1 (1578661-C1). Images were acquired using a Zeiss LSM880 confocal microscope and analyzed in Fiji. To quantify gene expression within the hair follicle, the freehand selection function was used to circle the hair bulb region on the max projection image, and the average fluorescent intensity was calculated using the *Measure* function.

### Growth plate RNA sequencing

To characterize gene expression during tail growth, we performed RNA-sequencing of congenic mouse vertebral growth plates (6 *std/std*, 6 *std/inv*, 5 *inv/inv* mice). We dissected growth plates from caudal vertebrae 8-10 of P25 mice. Growth plates were homogenized using the TissueLyser II (Qiagen) and then RNA was extracted using the Direct-zol RNA Miniprep Plus Kit (Zymo). RNA was quantified using the TapeStation (Agilent) and then RNA libraries were prepared using the Kapa mRNA Hyperprep kit. Quality control on RNA libraries was performed with the TapeStation (Agilent) and qPCR. RNA libraries were sequenced as 2x50bp or 2x150bp paired-end reads, across multiple NovaSeq SP Flow Cells (Illumina). We aligned the sequencing reads to the *P. maniculatus* transcriptome using STAR (*48*) and quantified transcript abundances using RSEM (*49*), as described above for melanocyte RNA sequencing. We identified differentially expressed genes using DESeq2 (*50*), comparing *inv/inv* v. *std/std* mice, with a significance threshold of 0.05 for the adjusted p-value and a log2FoldChange threshold of 1.

To test for allele-specific expression of *Npr3*, the top tail length candidate gene, we first identified haplotype-informative SNPs within *Npr3* UTRs or exons. We identified SNPs where all 6 *std/inv* samples were heterozygous at the SNP and one allele was fixed within one haplotype (*inv* or *std*), ensuring that the other allele came from the other haplotype (*inv* or *std*). We found a total of 12 haplotype-informative SNPs within *Npr3* and then recorded the number of RNA-sequencing reads with each allele in the 6 *std/inv* growth plate samples. Normalized allele count was calculated as the number of reads with one allele divided by the total number of reads at that SNP, for each sample.

We also performed RNA-sequencing on tibia growth plates for 3 *std/std* and 3 *inv/inv* mice. Tibia growth plates were dissected at P25. RNA extraction, sequencing and differential expression analyses were performed as described above.

### P. m. rufinus phenotypes and genome assembly

Coat color and tail length phenotypes were obtained for the *P. m. rufinus* population using museum specimens from the Museum of Southwestern Biology at the University of New Mexico (Table S2). Specimens were measured for coat color and tail length, as described above. We genotyped *P. m. rufinus* samples for the inversion using 2 Taqman SNP assays that differentiate the inversion and standard haplotypes within *P.m.rufinus*; these SNPs were selected based on whole-genome resequencing data from *P.m.rufinus* (see Methods section titled *Distribution of the inversion across populations*). For experimental analyses, *P.m.rufinus* mice were obtained from a colony in Dr. David Safronetz’s lab at the University of Manitoba. To create a *de novo* long-read genome assembly for *P. m. rufinus*, we selected a mouse homozygous for the inversion. We extracted high-molecular weight DNA from whole-blood using the Nanobind kit (PacBio), created a SMRTbell library using the SMRTbell Express Template Prep Kit 2.0 (PacBio), and sequenced using 5 SMRT cells on the Sequel II sequencer (PacBio). The genome assembly was created using *hifi-asm*, resulting in a contig-level genome assembly with contig N50 of 22.4 Mb. We then aligned the *P. m. rufinus* genome with the forest mouse genome (GenBank: GCA_049852335.1) using *minimap* v2.24 with *asm5*. *P. m. rufinus* contigs mapping to chr15 of the forest mouse genome were selected and synteny plots for chr15 were made in R (*52*).

### Fine-mapping of the tail length locus

We selected 2 *P.m.rufinus* males homozygous for the inversion allele (*inv^t^/inv^t^*) to breed with 2 congenic mouse females homozygous for the inversion (*inv^T^/inv^T^*). F1 hybrids were then intercrossed to create a mapping population of 373 F2 hybrids. Standard measurements of total length and tail length, as described above, were recorded for the F2 hybrids at ∼P90 (with ages ranging from P88 – P97). We performed low-coverage whole-genome sequencing (∼0.5x coverage) of the F2 hybrids, and high-coverage whole-genome sequencing (∼15x coverage) of the founding (F0) *rufinus* (n = 2) and congenic forest (n = 2) mice. Genomic DNA extraction was carried out as described above. Input DNA was quantified with the TapeStation (Agilent) and then DNA libraries were prepared using the Illumina DNA library preparation kit at 1⁄4 volume. Quality control on DNA libraries was performed with the TapeStation (Agilent) and qPCR. The libraries were then sequenced as 2x150bp paired-end reads, across two NovaSeq SP Flow Cells (Illumina). We mapped all samples to the *P. maniculatus* reference genome using bwa-mem. For the 4 founder mice, we performed joint genotyping with a large cohort of *P. maniculatus* mice, using GATK. For each breeding pair of *rufinus* x forest founders, we then selected SNPs that were opposite homozygous genotypes between the two mice; these SNPs allow for ancestry determination across the genomes of F2 hybrids. We next genotyped the F2 hybrids at these sets of ancestry-informative SNPs, using the SNPs corresponding to each F2’s founding breeder pair. To do so, we extracted the set of SNPs from the F2 bam files using *pileup* (samtools). We then ran the *fitHMM* step of MSG (*53*) to determine local ancestry. We extracted local average ancestry probabilities for each 100-kb window across the genome, and then imported the ancestry calls and phenotypes into r/qtl (*54*). We performed QTL mapping of tail length using marker-regression in r/qtl. Note that we re-arranged markers on chromosome 15 relative to the reference genome, such that the ordering of markers reflects the *inv* haplotype (instead of the *std* haplotype). We determined a genome-wide significance threshold using 1,000 permutations and a confidence interval on the chr15 QTL using a LOD-1 interval. Percent variance explained for each QTL was obtained using the formula 1 – 10^-(2/n)*LOD^.

### Distribution of the inversion across populations

To investigate the history of the inversion’s establishment and association with coat color and tail length, we first analyzed the distribution of the inversion across the species range using previously published data (*15, 16*). In addition, we included *P. m. rufinus* samples (population d) from the Museum of Southwestern Biology (University of New Mexico) and *P. m. rubidus* (population b) and *P. m. artemisiae* (population c) samples from the Museum of Comparative Zoology (Harvard University) (Table S2). We used Taqman SNP assays to genotype these samples for the inversion. To ensure that the SNPs were fixed between the inversion and standard haplotypes in each population, we analyzed whole-genome sequencing data for *n*=17, 17, 11, 15, 21 samples for the forest ecotype, prairie ecotype and populations b, c, d respectively (Table S2). We genotyped these samples for the inversion using principal component analyses of SNPs from the inversion region in *scikit-allel*; the top principal component separates samples by inversion genotype. We confirmed that the Taqman SNP assays selected for each population perfectly segregated with inversion genotypes in that population (Figure S11).

We next phenotyped a subset of populations for tail length and coat color. We selected populations in which the inversion was segregating at intermediate frequencies, and for which we had access to specimens, which yielded four populations (a,b,c,d). For mice from each population, we measured tail length and coat color as described above. We tested for significant effects of the inversion on coat color using t-tests for each population. We tested for significant effects of the inversion on tail length using linear models correcting for body size (populations a and b) or sacrum length measured from X-ray images as a proxy for body size (populations c and d): tail length ∼ inversion genotype + body size. Because population b samples came from both mainland and island subpopulations, we additionally accounted for subpopulation in population b using a linear mixed effects model: tail length ∼ inversion genotype + body size + (1|subpopulation).

### Maximum likelihood trees

To create a maximum likelihood tree for the inversion region across populations, we used a previously generated whole-genome resequencing dataset (*15*) in which samples were sequenced at 10-15x coverage. We selected 3 mice per homozygous genotype (*inv/inv*; *std/std*) per population, except for population c which had only 1 *inv/inv* sequenced mouse. We included *Peromyscus leucopus* as an outgroup. We used RAxML to build a maximum-likelihood tree for the inversion region (chr15:0-40 Mb) (*55*). Using a previously generated vcf (*15*), we selected biallelic SNPs within the inversion region. We filtered for SNPs with at least 60% of samples genotyped and thinned to 1 SNP per 100bp. We converted the vcf to a PHYLIP matrix using vcf2phylip.py (https://github.com/edgardomortiz/vcf2phylip) and removed invariant sites using ascbias.py (https://github.com/btmartin721/raxml_ascbias), resulting in a total of 48,487 SNPs. We then ran RAxML (v8.2.12) with the ASC_GTRCAT model and --asc-cor=lewis to correct for ascertainment bias associated with using SNPs. We performed 100 bootstraps and visualized the resulting tree in iTOL. We next performed a gene tree analysis using windowed maximum likelihood trees. We used RAxML as described above to create maximum likelihood trees for 500-kb non-overlapping windows across the inversion region, with 532-769 SNPs per window. We additionally created trees for 100-kb non-overlapping windows in the *Npr3* genomic region from chr15:22.5-23.5 Mb, with 104-146 SNPs per window. We then manually categorized trees by topology for each window.

---

**Figure S1.**
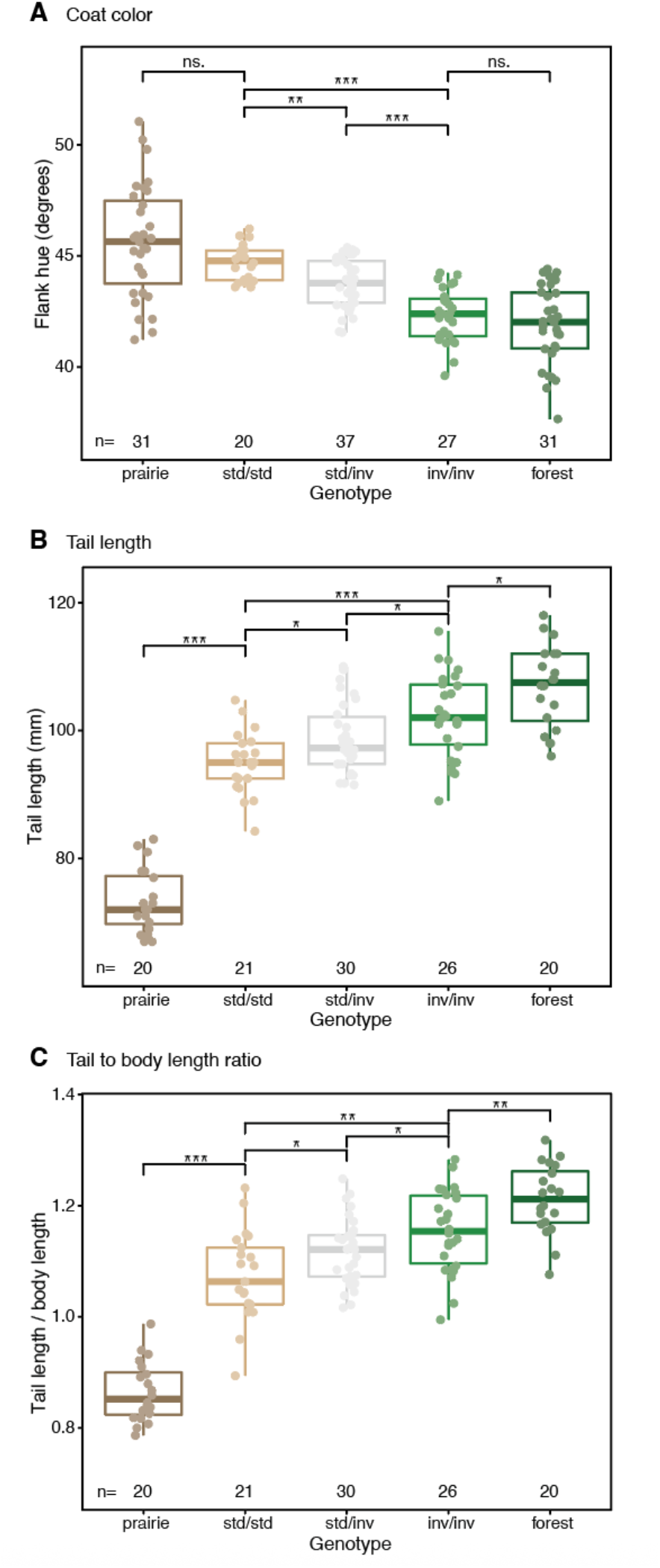
Tail length and coat color phenotypes in congenic mice, compared with forest and prairie ecotypes. (**A**) Coat color measured as flank hue (degrees) of adult mouse coat. (**B**) Tail length measurements in adult mice. (**C**) Ratio of tail length normalized by body length in adult mice. Symbols: ns. = *P* > 0.1; \**P* < 0.1; \*\**P* < 0.01; \*\*\**P* < 0.001 (two-sided Welch’s t-tests). Boxplots indicate the median (center line), the 25th and 75th percentiles (box extent); whiskers show largest or smallest value within 1.5 times the inter-quartile range; dots show individual data points.

**Figure S2.**
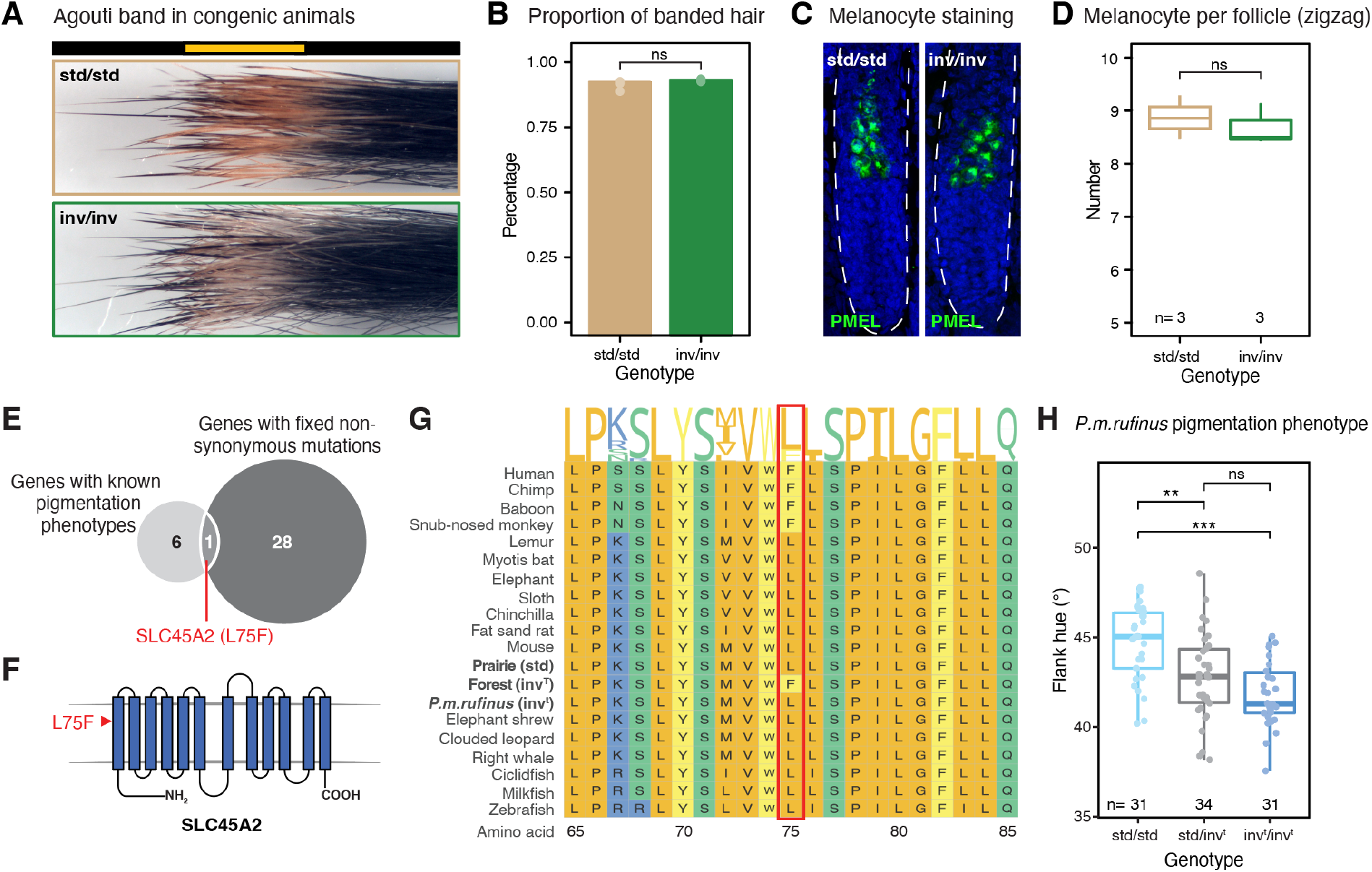
Coat color effects of the inversion are unlikely mediated through coding changes. (**A**) Banding analysis of hair bundles collected from flanks of adult *std/std* and *inv/inv* mice. (**B**) Proportions of banded hair between genotypes. Sample sizes: 3 *std/std* mice and 3 *inv/inv* mice, with 200 hairs per animal analyzed. No statistical significance found (t-test, *P* > 0.05). (**C**) Representative images of hair follicles stained with an antibody against PMEL, showing properly specified melanocytes in both genotypes. (**D**) Quantification of PMEL^+^ melanocytes per hair follicle between genotypes. No statistical significance found (paired t-test, *P* > 0.05). (**E**) Overlap between genes within the inversion with known pigmentation phenotypes (n=7) and genes within the inversion with fixed non-synonymous mutations (n=29). Only 1 gene, *Slc45a2*, is found in this intersection. (**F**) Structure of SLC45A2, which contains 12 transmembrane domains, with the L75F coding mutation found between the *std* and *inv* haplotypes highlighted in red. (**G**) Conservation of SLC45A2 amino acids across different mammalian species. The deer mouse sequences are highlighted in bold. (**H**) Coat color phenotypes, measured as flank hue (degrees) of *P. m. rufinus* specimens with different inversion genotypes. Symbols: ns. = *P* > 0.1; \*\**P* < 0.01; \*\*\**P* < 0.001 (two-sided Welch’s t-tests). Boxplots indicate the median (center line), the 25th and 75th percentiles (box extent); whiskers show largest or smallest value within 1.5 times the inter-quartile range; dots show individual data points.

**Figure S3.**
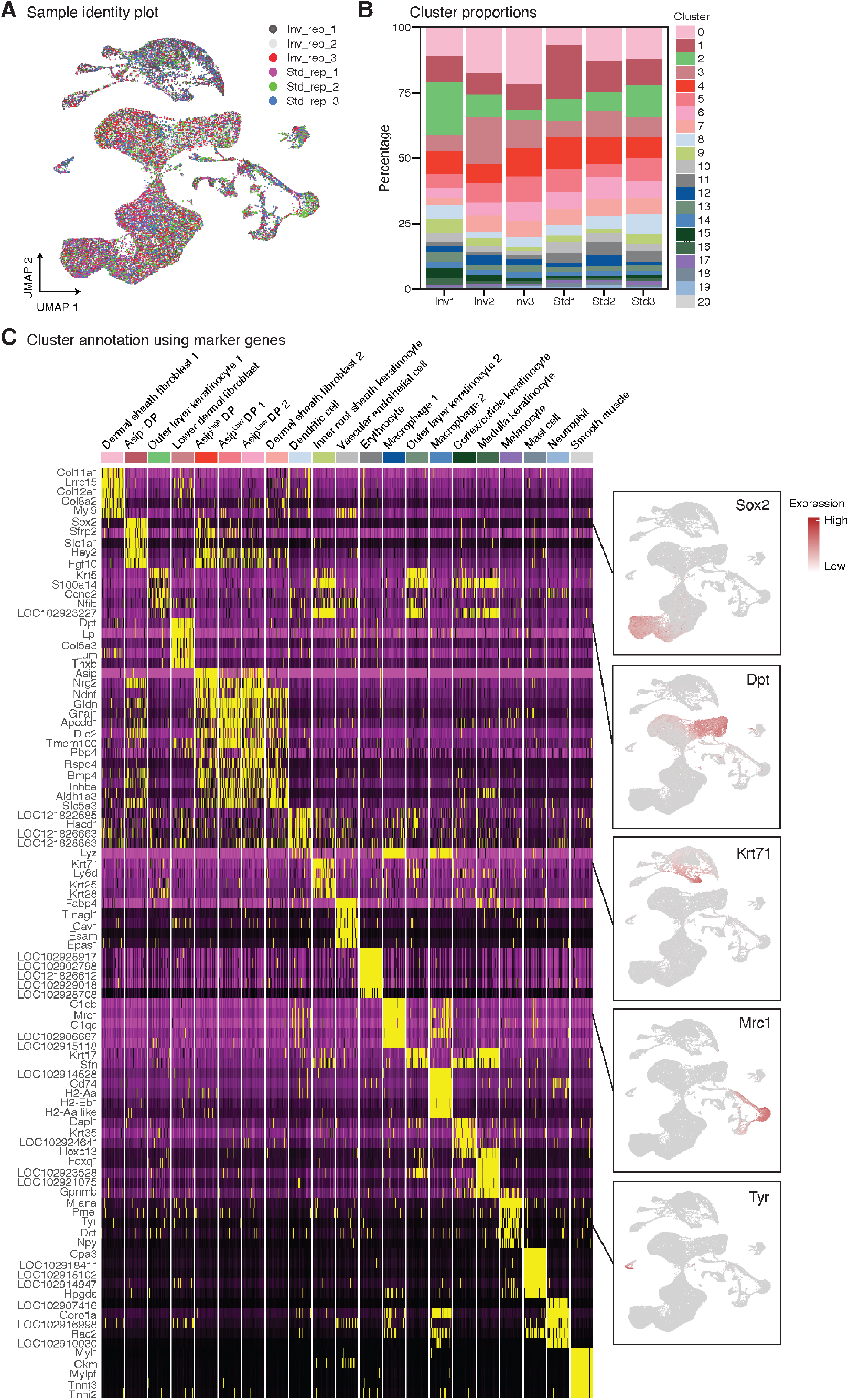
Single-cell sequencing of anagen skin from congenic deer mice. (**A**) UMAP plot colored by mouse sample IDs, showing even distribution of samples across clusters. (**B**) Proportions of cell clusters per sample. (**C**) Heatmap showing the expression of top 5 markers for each cluster. Cluster identities were assigned based on previously published cell-type markers in *Mus musculus* (*56*). Feature plots for key marker genes are shown on the right.

**Figure S4.**
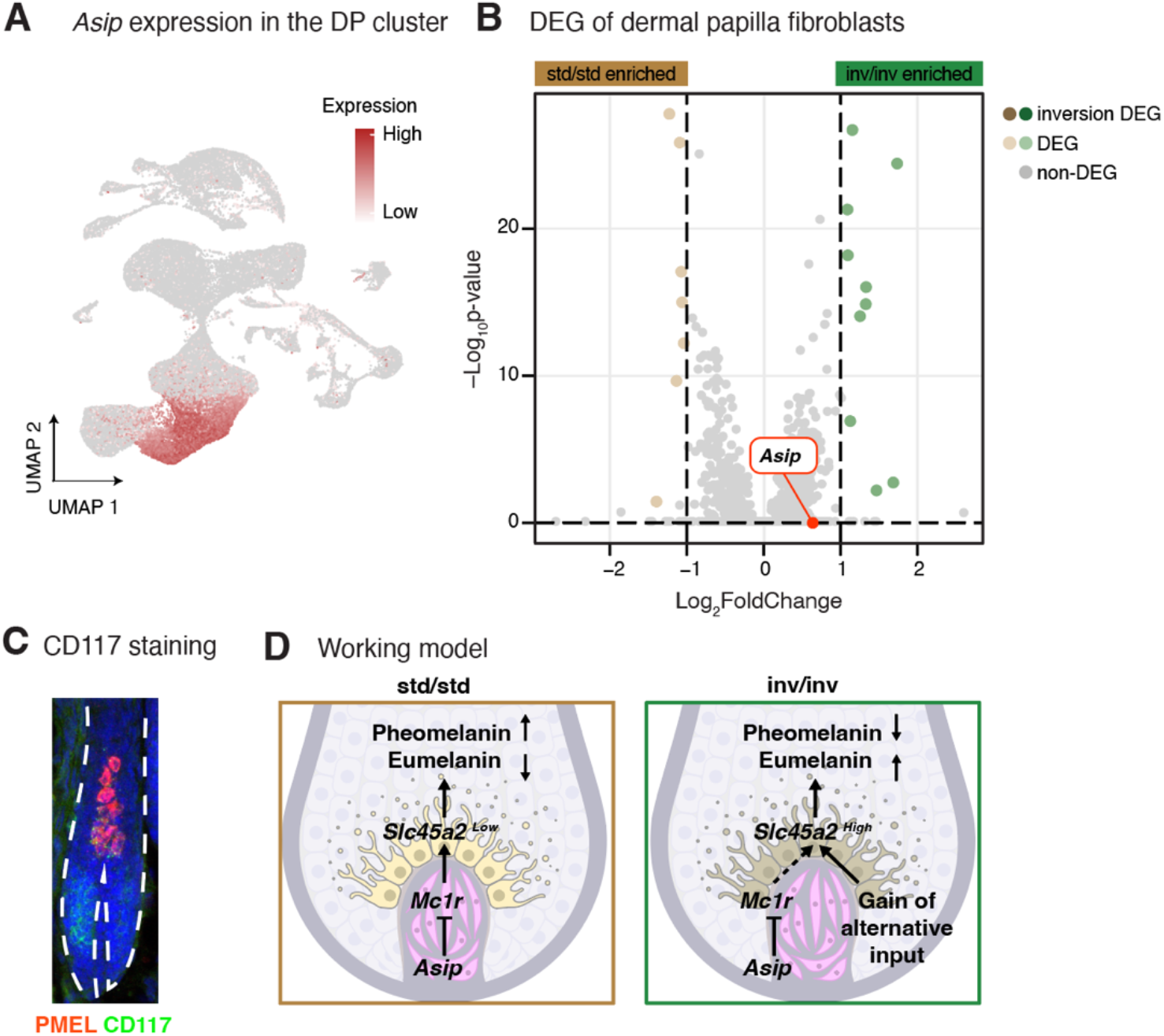
A working model for *Slc45a2*. (**A**) Feature plot of *Asip* showing high expression in the DP cluster. (**B**) DEG analysis of dermal papilla fibroblasts identified from single cell RNA-seq. No inversion genes are differentially expressed. *Asip* is highlighted in red. (**C**) Representative immunofluorescence image of CD117 and PMEL in deer mouse anagen hair follicle. (**D**) A working model explaining how gain of alternative regulatory input in the *inv* allele alleviates *Slc45a2* from transcriptional repression of *Asip*.

**Figure S5.**
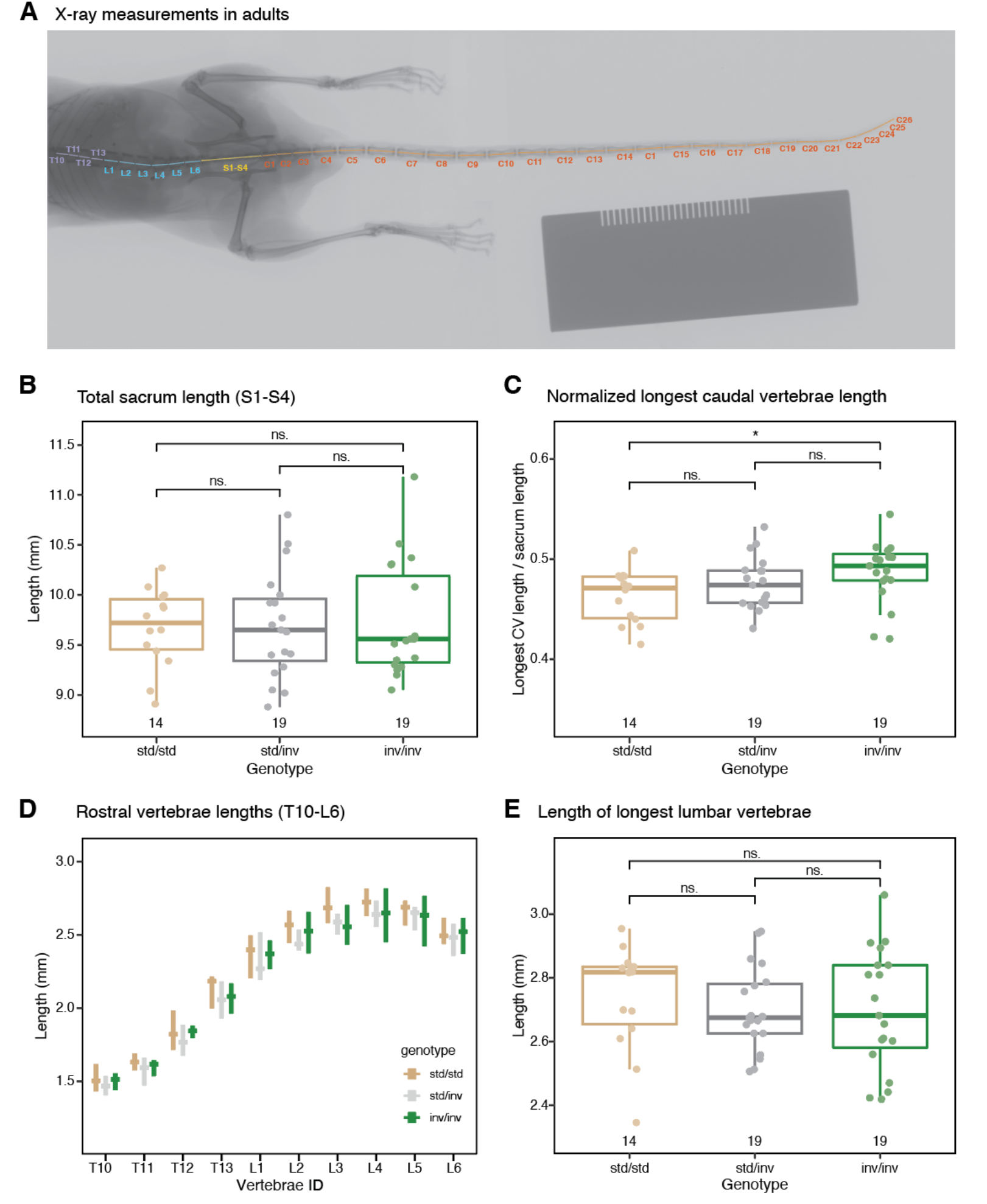
Vertebral measurements. (**A**) Labeled X-ray image of an adult deer mouse, demonstrating the positions of rostral thoracic vertebrae (purple, T10-T13), lumbar vertebrae (blue, L1 – L6), sacral vertebrae (yellow, S1 – S4), and caudal vertebrae (orange, C1 – C26) whose lengths were measured in the following panels. (**B** – **E**) Measurements from X-ray images of congenic mice at P70-75. (**B**) Length of the sacrum, by summing the S1 to S4 vertebrae. (**C**) Ratio of the length of the longest caudal vertebrae to the length of the sacrum. (**D**) Lengths of rostral thoracic and lumbar vertebrae. (**E**) Length of the longest lumbar vertebra. Symbols: ns. = *P* > 0.05; \**P* < 0.05 (two-sided Welch’s t-tests). Boxplots indicate the median (center line), the 25th and 75th percentiles (box extent); whiskers show largest or smallest value within 1.5 times the inter-quartile range; dots show individual data points.

**Figure S6.**
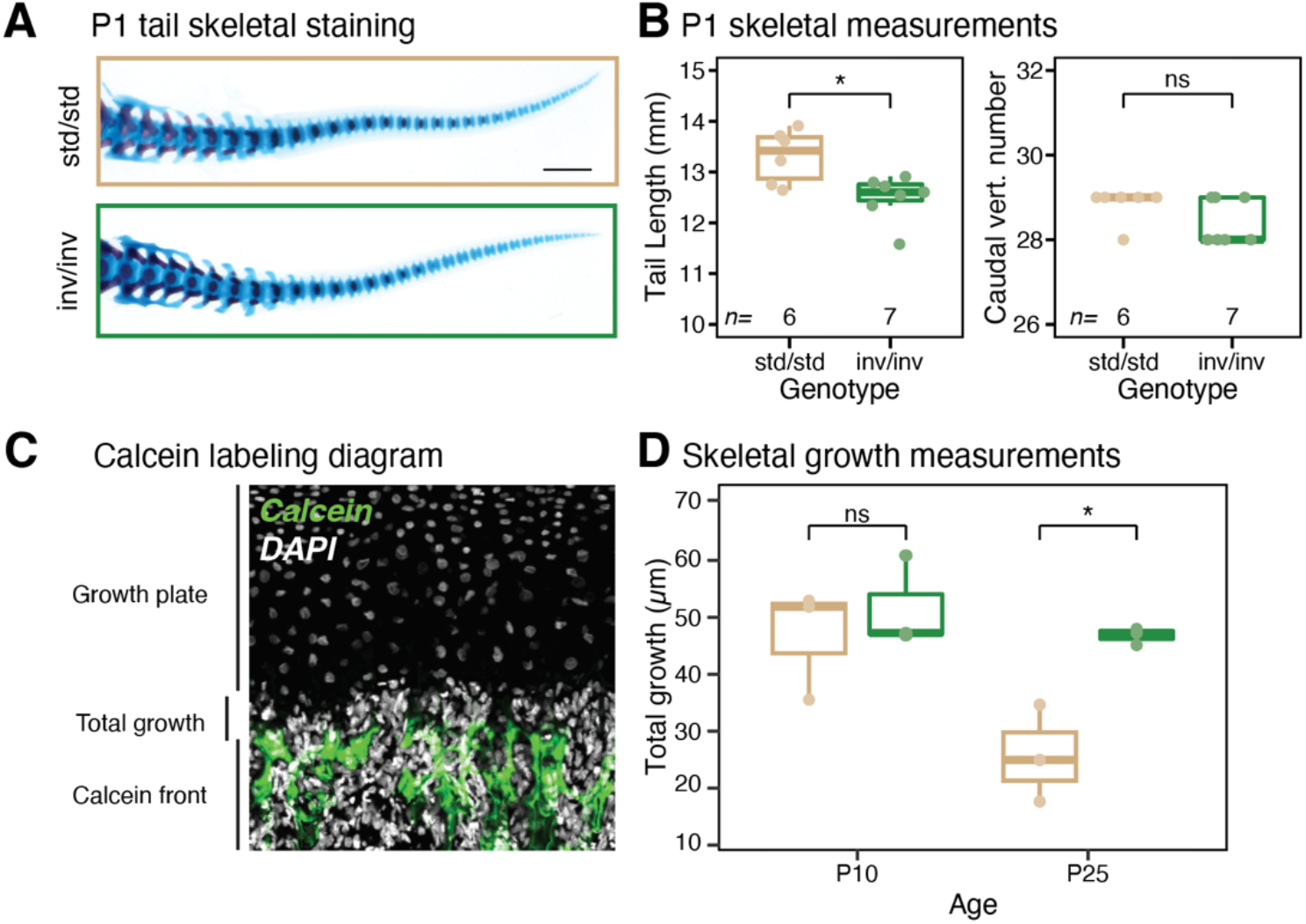
Vertebral growth measurements. (**A**) Representative skeletal staining of *std/std* and *inv/inv* P1 pups. Tail length and caudal vertebral number are quantified in (**B**). Symbols: ns. = *P* > 0.05; \**P* < 0.05 (two-sided Welch’s t-tests). (**C**) Diagram of the calcein pulse-chase experiment with an example image of a caudal vertebral growth plate. Cumulative growth over three days is quantified in (**D**). Symbols: ns. = *P* > 0.05; \**P* < 0.05 (two-sided Welch’s t-tests).

**Figure S7.**
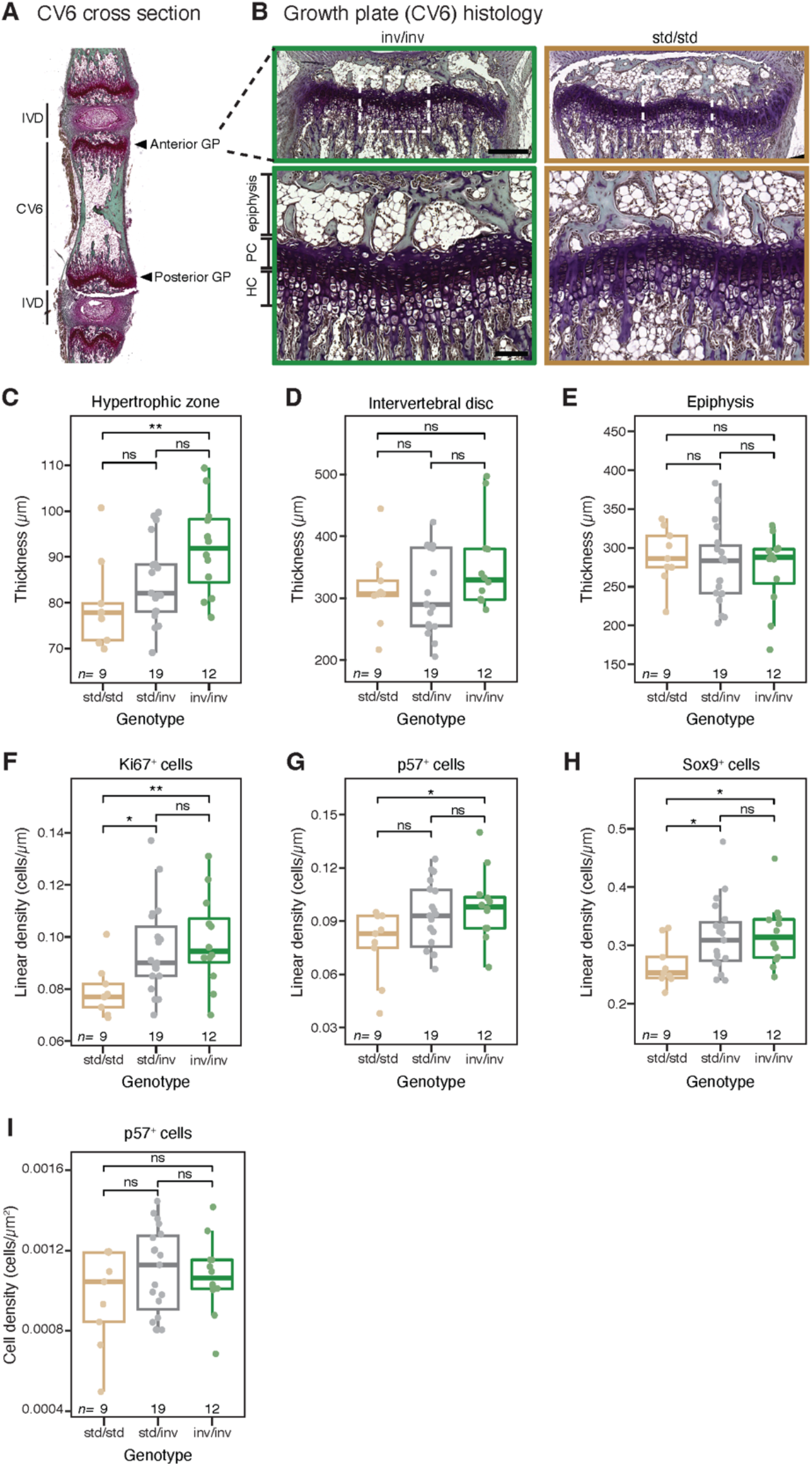
CV_6_ growth plate histology. (**A**) A representative cross section of CV_6_, with anterior and posterior intervertebral discs (IVD) and growth plates (GP) highlighted. (**B**) Representative histology images for the anterior growth plates of CV_6_ in *inv/inv* and *std/std* mice. Insets below show zoomed-in regions outlined by the white box above. The epiphysis, proliferating chondrocyte zone (PC), and hypertrophic chondrocyte zone (HC), are labeled. (**C**) Quantification of hypertrophic chondrocyte zone thickness. (**D**) Quantification of intervertebral disc thickness. (**E**) Quantification of epiphyseal thickness. (**F**) Linear density of Ki67^+^ cells which marks proliferative chondrocytes (total Ki67^+^ cells/diameter of the growth plate). (**G**) Linear density of p57^+^ cells which marks hypertrophic chondrocytes (total p57^+^ cells/ diameter of the growth plate). (**H**) Linear density of SOX9^+^ cells which marks the entire growth plate (total SOX9^+^ cells/ diameter of the growth plate). (**I**) Quantification of hypertrophic chondrocyte density, marked by p57. Symbols: ns. = *P* > 0.05; \**P* < 0.05 (two-sided Welch’s t-tests). Boxplots indicate the median (center line), the 25th and 75th percentiles (box extent); whiskers show largest or smallest value within 1.5 times the inter-quartile range; dots show individual data points.

**Figure S8.**
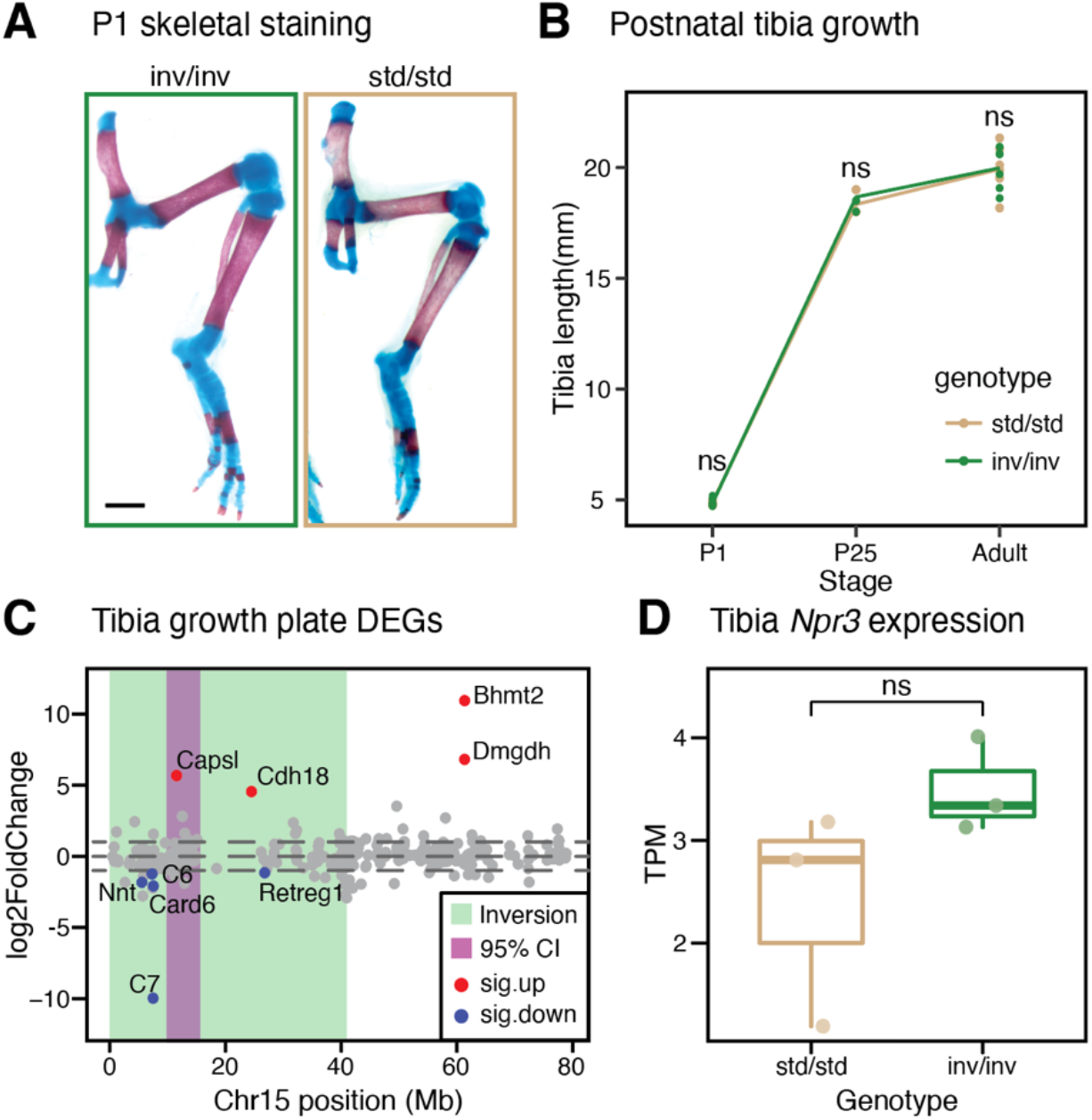
Measurements of tibia lengths and gene expression in congenic deer mice. (**A**) Representative images of P1 *inv/inv* and *std/std* tibia. (**B**) Post-natal tibia growth rates measured at P1, P25 and adult (P70-75) for *inv/inv* and *std/std* mice; no significant differences in tibia lengths were found. (**C**) Differentially expressed genes between *inv/inv* and *std/std* P25 tibia growth plates along chromosome 15. X-axis shows gene positions and y-axis shows log2FoldChange. The inversion region is colored in green, with the fine-mapping QTL highlighted in purple. Significant DEGs are highlighted in red (higher expression in *std/std* mice) or blue (lower expression in *std/std* mice). (**D**) Expression of *Npr3* in tibia growth plates from 3 *std/std* and 3 *inv/inv* mice. TPM=transcripts per million. Symbols: ns = *P* > 0.05 (Welch’s two-sided t-test).

**Figure S9.**
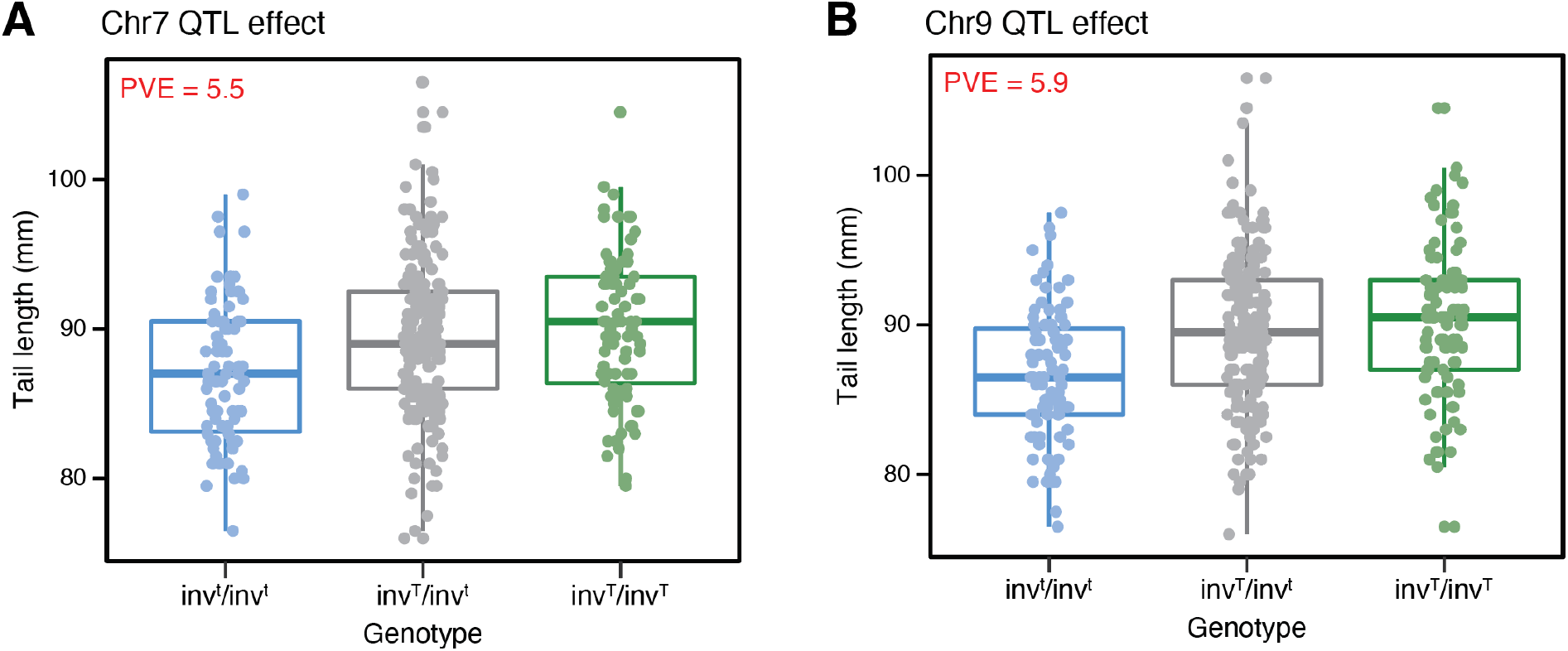
The effects of additional fine-mapping QTLs on tail length. Effects of the chr7 QTL. (**A**) and chr9 QTL (**B**) in the F2 hybrids, with F2s grouped by genotype at the peak marker (*inv^t^* = *P. m. rufinus* allele, *inv^T^* = forest allele). PVE = percent variance explained.

**Figure S10.**
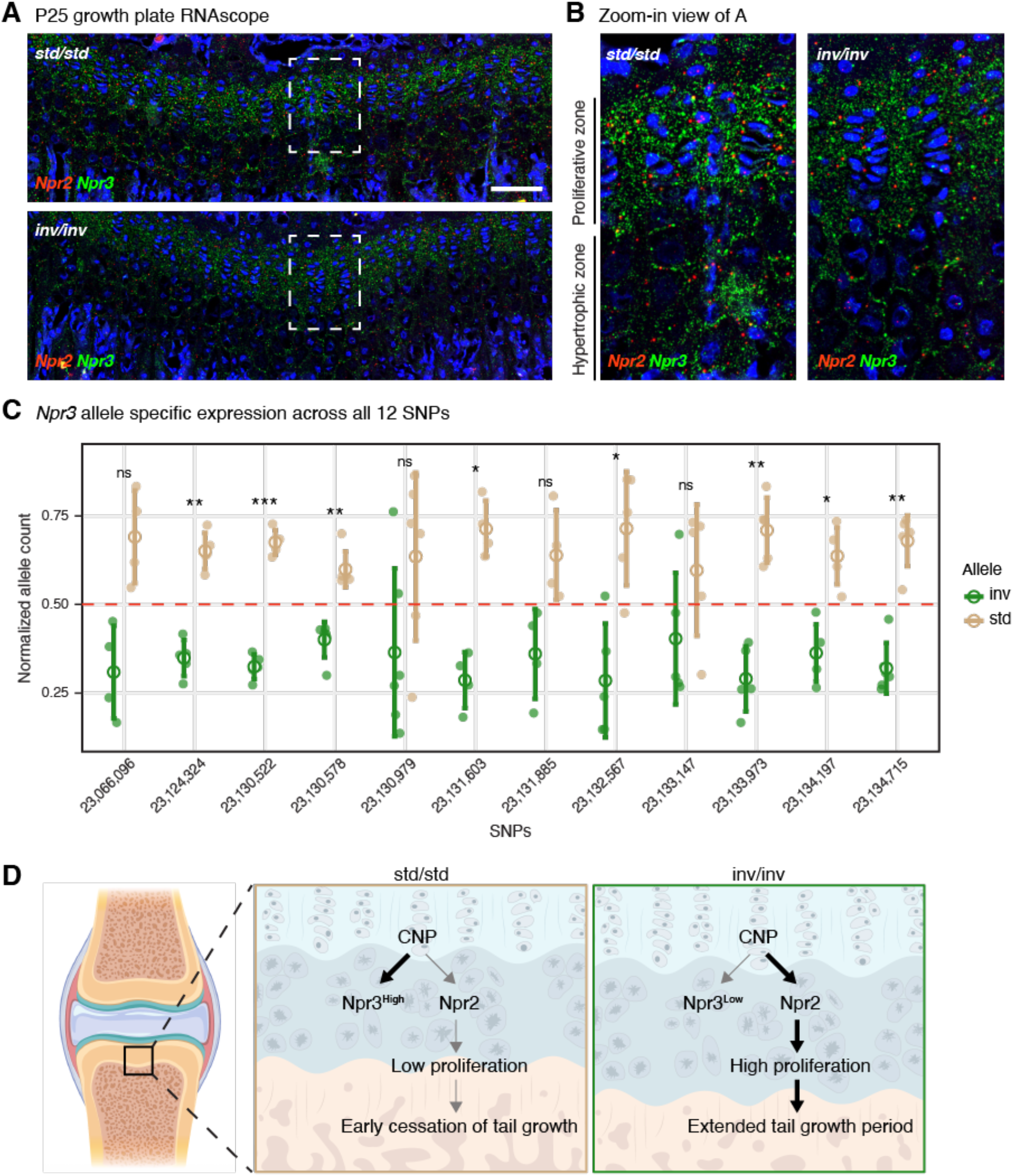
Allele-specific expression of *Npr3* underlies tail length difference. (**A**) Representative images of CV_6_ growth plates showing mRNAs of *Npr3* (green) and *Npr2* (red) detected by RNAscope *in situ*. White boxes indicate zoomed-in regions shown in (**B**). (**C**) Allele-specific expression of 12 informative SNPs within *Npr3* exons. SNP locations are labeled on the x-axis, with normalized allele count (number of reads from a specific allele/total number of reads at this SNP) shown on the y-axis. Open circles show the average value, with vertical lines indicating standard deviations. Dashed red line indicates expected normalized allele count with no allele-specific expression. Dots are individual samples. Symbols: ns. = *P* > 0.05; \**P* < 0.05; \*\**P* < 0.01; \*\*\**P* < 0.001 (paired t-tests). (**D**) Model of how *Npr3* differential expression mediates tail length. In *std/std* mice, *Npr3* expression is high in CV growth plates at P25, leading to reduced *Npr2* activity and thereby reduced proliferation of chondrocytes, resulting in an earlier cessation of tail growth. In *inv/inv* mice, *Npr3* expression is low, leading to increased *Npr2* activity and thereby increased proliferation of chondrocytes, resulting in a prolonged tail growth period.

**Figure S11.**
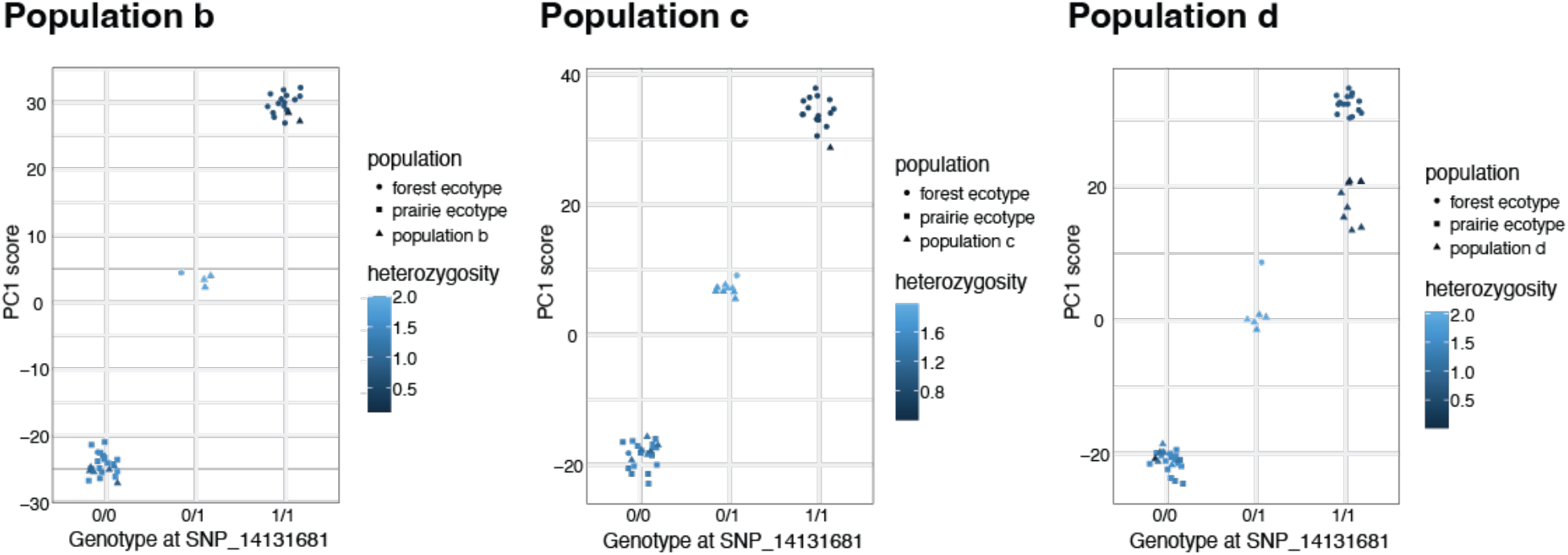
Genotyping mouse populations for the inversion using SNP assays. Taqman assay results for an example SNP used for genotyping the inversion across populations. For populations b-d, genotypes at this SNP (chr15:14,131,681) are shown along the x-axis, with PC1 score of the inversion region determined by whole-genome sequencing shown along the y-axis. Points are colored by average heterozygosity across the inversion region. SNP genotypes perfectly correlate with inversion genotypes inferred from whole-genome sequencing using PCA. Data from the original forest (indicated by circles) and prairie (indicated by squares) populations are also included in each plot as references.

**Table S1.**
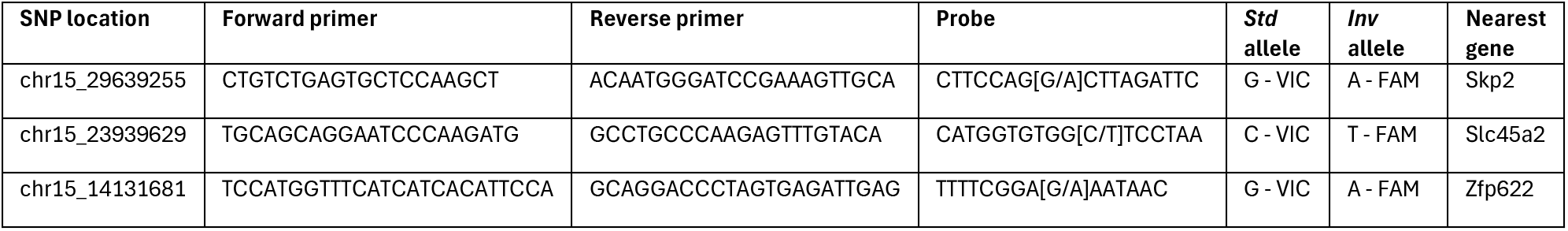
Taqman SNP assays for genotyping the inversion. Three Taqman SNP assays used for genotyping the inversion across populations. SNP location, sequences for the forward primer, reverse primer, and probe, the *std* and *inv* alleles, and the nearest gene are provided.

**Table S2.** Specimen IDs for population sampling. The IDs for deer mouse specimens genotyped for the inversion. Population (pop), study-specific ID (ID), museum specimen ID (Museum ID), museum, inversion genotype and NCBI accession number are provided (unassigned will be assigned upon uploading to NCBI SRA).

| Pop | ID | Museum ID | Museum | Inversion genotype | NCBI accession |
| --- | --- | --- | --- | --- | --- |
| a | ERH88 | 70288 | HU_MCZ | std/std | unassigned |
| a | ERH89 | 70289 | HU_MCZ | std/std | unassigned |
| a | ERH107 | 70300 | HU_MCZ | std/std | unassigned |
| a | ERH90 | 70290 | HU_MCZ | std/std | unassigned |
| a | ERH94 | 69534 | HU_MCZ | std/std | unassigned |
| a | ERH83 | 70286 | HU_MCZ | std/std | unassigned |
| a | ERH108 | 69536 | HU_MCZ | std/std | unassigned |
| a | ERH87 | 70287 | HU_MCZ | std/std | unassigned |
| a | ERH110 | 70302 | HU_MCZ | std/std | unassigned |
| a | ERH93 | 70291 | HU_MCZ | std/std | unassigned |
| a | ERH95 | 70292 | HU_MCZ | std/std | unassigned |
| a | ERH100 | 70297 | HU_MCZ | std/std | unassigned |
| a | ERH99 | 70296 | HU_MCZ | std/std | unassigned |
| a | ERH97 | 70294 | HU_MCZ | std/std | unassigned |
| a | ERH96 | 70293 | HU_MCZ | std/std | unassigned |
| a | ERH139 | 70330 | HU_MCZ | inv/inv | unassigned |
| a | ERH136 | 70327 | HU_MCZ | inv/inv | unassigned |
| a | ERH141 | 70332 | HU_MCZ | inv/inv | unassigned |
| a | ERH143 | 70334 | HU_MCZ | inv/inv | unassigned |
| a | ERH121 | 70312 | HU_MCZ | inv/inv | unassigned |
| a | ERH120 | 70311 | HU_MCZ | std/std | unassigned |
| a | ERH125 | 70316 | HU_MCZ | std/std | unassigned |
| a | ERH135 | 70326 | HU_MCZ | std/std | unassigned |
| a | ERH145 | 70336 | HU_MCZ | inv/inv | unassigned |
| a | ERH144 | 70335 | HU_MCZ | inv/inv | unassigned |
| a | ERH150 | 70341 | HU_MCZ | inv/inv | unassigned |
| a | ERH149 | 70340 | HU_MCZ | inv/inv | unassigned |
| a | ERH161 | 70352 | HU_MCZ | inv/inv | unassigned |
| a | ERH163 | 70354 | HU_MCZ | inv/inv | unassigned |
| a | ERH157 | 70348 | HU_MCZ | std/std | unassigned |
| a | ERH156 | 70347 | HU_MCZ | inv/inv | unassigned |
| a | ERH166 | 70357 | HU_MCZ | inv/inv | unassigned |
| a | ERH167 | 70358 | HU_MCZ | inv/inv | unassigned |
| a | ERH172 | 70363 | HU_MCZ | inv/inv | unassigned |
| a | ERH177 | 70368 | HU_MCZ | inv/inv | unassigned |
| a | ERH183 | 70374 | HU_MCZ | inv/inv | unassigned |
| a | ERH174 | 70365 | HU_MCZ | inv/inv | unassigned |
| a | ERH179 | 70370 | HU_MCZ | inv/inv | unassigned |
| a | ERH178 | 70369 | HU_MCZ | inv/inv | unassigned |
| a | ERH186 | 70377 | HU_MCZ | inv/inv | unassigned |
| a | ERH189 | 70380 | HU_MCZ | inv/inv | unassigned |
| a | ERH205 | 70396 | HU_MCZ | inv/inv | unassigned |
| a | ERH187 | 70378 | HU_MCZ | inv/inv | unassigned |
| a | ERH199 | 70390 | HU_MCZ | inv/inv | unassigned |
| a | ERH202 | 70393 | HU_MCZ | inv/inv | unassigned |
| a | ERH200 | 70391 | HU_MCZ | inv/inv | unassigned |
| a | ERH191 | 70382 | HU_MCZ | inv/inv | unassigned |
| a | ERH193 | 70384 | HU_MCZ | inv/inv | unassigned |
| a | ERH195 | 70386 | HU_MCZ | inv/inv | unassigned |
| a | ERH217 | 70408 | HU_MCZ | inv/inv | unassigned |
| a | ERH194 | 70385 | HU_MCZ | inv/inv | unassigned |
| a | ERH218 | 70409 | HU_MCZ | inv/inv | unassigned |
| a | ERH210 | 70401 | HU_MCZ | inv/inv | unassigned |
| a | ERH214 | 70405 | HU_MCZ | inv/inv | unassigned |
| a | ERH188 | 70379 | HU_MCZ | inv/inv | unassigned |
| a | ERH190 | 70381 | HU_MCZ | inv/inv | unassigned |
| a | ERH215 | 70406 | HU_MCZ | inv/inv | unassigned |
| a | ERH213 | 70404 | HU_MCZ | inv/inv | unassigned |
| a | ERH204 | 70395 | HU_MCZ | inv/inv | unassigned |
| a | ERH222 | 70413 | HU_MCZ | std/std | unassigned |
| a | ERH212 | 70403 | HU_MCZ | inv/inv | unassigned |
| a | ERH55 | 70261 | HU_MCZ | inv/inv | unassigned |
| a | ERH62 | 70268 | HU_MCZ | inv/inv | unassigned |
| a | ERH60 | 70266 | HU_MCZ | inv/inv | unassigned |
| a | ERH61 | 70267 | HU_MCZ | inv/inv | unassigned |
| a | ERH56 | 70262 | HU_MCZ | inv/inv | unassigned |
| a | ERH59 | 70265 | HU_MCZ | inv/inv | unassigned |
| a | ERH71 | 70277 | HU_MCZ | inv/inv | unassigned |
| a | ERH58 | 70264 | HU_MCZ | inv/inv | unassigned |
| a | ERH57 | 70263 | HU_MCZ | inv/inv | unassigned |
| a | ERH66 | 70272 | HU_MCZ | inv/inv | unassigned |
| a | ERH65 | 70271 | HU_MCZ | inv/inv | unassigned |
| a | ERH74 | 70280 | HU_MCZ | inv/inv | unassigned |
| a | ERH67 | 70273 | HU_MCZ | inv/inv | unassigned |
| a | ERH68 | 70274 | HU_MCZ | inv/inv | unassigned |
| a | ERH75 | 70281 | HU_MCZ | inv/inv | unassigned |
| a | ERH70 | 70276 | HU_MCZ | inv/inv | unassigned |
| a | ERH69 | 70275 | HU_MCZ | inv/inv | unassigned |
| a | ERH79 | 70285 | HU_MCZ | inv/inv | unassigned |
| a | ERH26 | 70248 | HU_MCZ | std/std | unassigned |
| a | ERH12 | 70240 | HU_MCZ | inv/inv | unassigned |
| a | ERH27 | 69544 | HU_MCZ | inv/inv | unassigned |
| a | ERH14 | 70242 | HU_MCZ | inv/inv | unassigned |
| a | ERH42 | 70254 | HU_MCZ | inv/inv | unassigned |
| a | ERH19 | 70244 | HU_MCZ | inv/inv | unassigned |
| a | ERH53 | 69556 | HU_MCZ | inv/inv | unassigned |
| a | ERH52 | 69555 | HU_MCZ | inv/inv | unassigned |
| a | ERH25 | 70247 | HU_MCZ | inv/inv | unassigned |
| a | ERH46 | 69562 | HU_MCZ | inv/inv | unassigned |
| a | ERH36 | 69566 | HU_MCZ | inv/inv | unassigned |
| a | ERH38 | 69558 | HU_MCZ | inv/inv | unassigned |
| a | ERH48 | 69563 | HU_MCZ | inv/inv | unassigned |
| b | FSB2 | 69714 | HU_MCZ | std/std | unassigned |
| b | FSB110 | 69726 | HU_MCZ | inv/inv | unassigned |
| b | FSB116 | 69727 | HU_MCZ | std/std | unassigned |
| b | FSB62 | 69728 | HU_MCZ | std/std | unassigned |
| b | FSB6 | 69733 | HU_MCZ | inv/inv | unassigned |
| b | FSB7 | 69734 | HU_MCZ | std/std | unassigned |
| b | FSB11 | 69737 | HU_MCZ | std/std | unassigned |
| b | FSB26 | 69744 | HU_MCZ | std/std | unassigned |
| b | FSB27 | 69745 | HU_MCZ | std/std | unassigned |
| b | FSB37 | 69752 | HU_MCZ | inv/inv | unassigned |
| b | FSB48 | 69756 | HU_MCZ | inv/inv | unassigned |
| b | FSB65 | 69761 | HU_MCZ | std/std | unassigned |
| b | FSB73 | 69765 | HU_MCZ | inv/inv | unassigned |
| b | FSB79 | 69767 | HU_MCZ | std/std | unassigned |
| b | FSB83 | 69770 | HU_MCZ | inv/inv | unassigned |
| b | FSB111 | 69778 | HU_MCZ | std/std | unassigned |
| b | FSB113 | 69779 | HU_MCZ | std/std | unassigned |
| b | FSB25 | 69784 | HU_MCZ | std/std | unassigned |
| b | FSB74 | 69785 | HU_MCZ | inv/inv | unassigned |
| b | FSB13 | 69789 | HU_MCZ | std/std | unassigned |
| b | FSB42 | 69793 | HU_MCZ | inv/inv | unassigned |
| b | FSB43 | 69794 | HU_MCZ | inv/inv | unassigned |
| b | FSB50 | 69796 | HU_MCZ | std/std | unassigned |
| b | FSB52 | 69797 | HU_MCZ | std/std | unassigned |
| b | FSB57 | 69799 | HU_MCZ | std/std | unassigned |
| b | FSB76 | 69803 | HU_MCZ | inv/inv | unassigned |
| b | FSB77 | 69804 | HU_MCZ | inv/inv | unassigned |
| b | FSB85 | 69807 | HU_MCZ | std/std | unassigned |
| b | FSB87 | 69809 | HU_MCZ | inv/inv | unassigned |
| b | FSB92 | 69812 | HU_MCZ | std/std | unassigned |
| c | 33296 | NA | HU_MCZ | std/std | unassigned |
| c | 33297 | NA | HU_MCZ | inv/inv | unassigned |
| c | 33298 | NA | HU_MCZ | inv/inv | unassigned |
| c | 33314 | NA | HU_MCZ | std/std | unassigned |
| c | 33394 | NA | HU_MCZ | inv/inv | unassigned |
| c | 33395 | NA | HU_MCZ | std/std | unassigned |
| c | 33396 | NA | HU_MCZ | inv/inv | unassigned |
| c | 33397 | NA | HU_MCZ | std/std | unassigned |
| c | 33398 | NA | HU_MCZ | std/std | unassigned |
| c | 33399 | NA | HU_MCZ | std/std | unassigned |
| c | 33400 | NA | HU_MCZ | std/std | unassigned |
| c | 33404 | NA | HU_MCZ | inv/inv | unassigned |
| c | 33406 | NA | HU_MCZ | std/std | unassigned |
| c | 33409 | NA | HU_MCZ | std/std | unassigned |
| c | 33410 | NA | HU_MCZ | std/std | unassigned |
| c | 33443 | NA | HU_MCZ | std/std | unassigned |
| c | 33444 | NA | HU_MCZ | inv/inv | unassigned |
| c | 33445 | NA | HU_MCZ | std/std | unassigned |
| c | 33446 | NA | HU_MCZ | inv/inv | unassigned |
| c | 33448 | NA | HU_MCZ | std/std | unassigned |
| c | 33449 | NA | HU_MCZ | std/std | unassigned |
| c | 33452 | NA | HU_MCZ | std/std | unassigned |
| c | 33453 | NA | HU_MCZ | std/std | unassigned |
| c | 33454 | NA | HU_MCZ | inv/inv | unassigned |
| d | 256610 | 280989 | UNM_MSB | inv/inv | unassigned |
| d | 256690 | 280947 | UNM_MSB | inv/inv | unassigned |
| d | 256691 | 280948 | UNM_MSB | std/std | unassigned |
| d | 256702 | 280959 | UNM_MSB | inv/inv | unassigned |
| d | 256704 | 280961 | UNM_MSB | inv/inv | unassigned |
| d | 256706 | 280963 | UNM_MSB | inv/inv | unassigned |
| d | 256707 | 280964 | UNM_MSB | std/std | unassigned |
| d | 256708 | 280965 | UNM_MSB | inv/inv | unassigned |
| d | 256709 | 280966 | UNM_MSB | inv/inv | unassigned |
| d | 256711 | 280968 | UNM_MSB | std/std | unassigned |
| d | 256718 | 280975 | UNM_MSB | std/std | unassigned |
| d | 256719 | 280976 | UNM_MSB | inv/inv | unassigned |
| d | 256720 | 280977 | UNM_MSB | inv/inv | unassigned |
| d | 256733 | 280697 | UNM_MSB | inv/inv | unassigned |
| d | 256736 | 280701 | UNM_MSB | inv/inv | unassigned |
| d | 256737 | 280702 | UNM_MSB | inv/inv | unassigned |
| d | 256741 | 280706 | UNM_MSB | std/std | unassigned |
| d | 256743 | 280708 | UNM_MSB | inv/inv | unassigned |
| d | 256746 | 280711 | UNM_MSB | inv/inv | unassigned |
| d | 256749 | 280714 | UNM_MSB | inv/inv | unassigned |
| d | 256755 | 280720 | UNM_MSB | inv/inv | unassigned |
| d | 256758 | 280723 | UNM_MSB | inv/inv | unassigned |
| d | 256763 | 280728 | UNM_MSB | inv/inv | unassigned |
| d | 256764 | 280729 | UNM_MSB | std/std | unassigned |
| d | 181791 | 262509 | UNM_MSB | std/std | unassigned |
| d | 181792 | 262510 | UNM_MSB | std/std | unassigned |
| d | 181794 | 262512 | UNM_MSB | inv/inv | unassigned |
| d | 181798 | 262516 | UNM_MSB | std/std | unassigned |
| d | 181799 | 262517 | UNM_MSB | inv/inv | unassigned |
| d | 181811 | 262527 | UNM_MSB | inv/inv | unassigned |
| d | 181812 | 262528 | UNM_MSB | inv/inv | unassigned |
| d | 181835 | 262552 | UNM_MSB | inv/inv | unassigned |
| d | 181840 | 262557 | UNM_MSB | inv/inv | unassigned |
| d | 181841 | 262558 | UNM_MSB | inv/inv | unassigned |
| d | 181842 | 262559 | UNM_MSB | inv/inv | unassigned |
| d | 257180 | 280637 | UNM_MSB | inv/inv | unassigned |
| d | 257183 | 280640 | UNM_MSB | inv/inv | unassigned |
| d | 257184 | 280641 | UNM_MSB | std/std | unassigned |
| d | 257190 | 280645 | UNM_MSB | std/std | unassigned |
| d | 257188 | 280647 | UNM_MSB | std/std | unassigned |
| d | 257189 | 280648 | UNM_MSB | std/std | unassigned |
| d | 256767 | 280732 | UNM_MSB | std/std | unassigned |
| d | 257095 | 284473 | UNM_MSB | std/std | unassigned |
| d | 257099 | 284497 | UNM_MSB | inv/inv | unassigned |
| d | 257108 | 284506 | UNM_MSB | std/std | unassigned |
| d | 257109 | 284507 | UNM_MSB | std/std | unassigned |
| d | 257114 | 284512 | UNM_MSB | std/std | unassigned |
| d | 257117 | 284515 | UNM_MSB | std/std | unassigned |
| d | 257123 | 284521 | UNM_MSB | std/std | unassigned |
| d | 257127 | 284525 | UNM_MSB | std/std | unassigned |
| d | 257132 | 284530 | UNM_MSB | std/std | unassigned |
| d | 257133 | 284531 | UNM_MSB | std/std | unassigned |
| d | 257140 | 284538 | UNM_MSB | std/std | unassigned |
| d | 257141 | 284539 | UNM_MSB | std/std | unassigned |
| d | 257152 | 284540 | UNM_MSB | std/std | unassigned |
| d | 257147 | 284545 | UNM_MSB | std/std | unassigned |
| d | NA | 284571 | UNM_MSB | std/std | unassigned |
| d | 257153 | 284576 | UNM_MSB | std/std | unassigned |
| d | 257155 | 284578 | UNM_MSB | std/std | unassigned |
| d | 257165 | 284588 | UNM_MSB | std/std | unassigned |
| d | NA | 284683 | UNM_MSB | inv/inv | unassigned |
| prairie | ERH108 | MCZ69536 | HU_MCZ | std/std | SRR13318593 |
| prairie | ERH87 | MCZ70287 | HU_MCZ | std/std | SRR13318653 |
| prairie | ERH99 | MCZ70296 | HU_MCZ | std/std | SRR13318667 |
| prairie | ERH95 | MCZ70292 | HU_MCZ | std/std | SRR13318700 |
| prairie | ERH97 | MCZ70294 | HU_MCZ | std/std | SRR13318678 |
| prairie | ERH100 | MCZ70297 | HU_MCZ | std/std | SRR13318615 |
| prairie | ERH110 | MCZ70302 | HU_MCZ | std/std | SRR13318725 |
| prairie | ERH83 | MCZ70286 | HU_MCZ | std/std | SRR13318714 |
| prairie | ERH88 | MCZ70288 | HU_MCZ | std/std | SRR13318642 |
| prairie | ERH89 | MCZ70289 | HU_MCZ | std/std | SRR13318631 |
| prairie | ERH90 | MCZ70290 | HU_MCZ | std/std | SRR13318758 |
| prairie | ERH96 | MCZ70293 | HU_MCZ | std/std | SRR13318689 |
| prairie | ERH107 | MCZ70300 | HU_MCZ | std/std | SRR13318604 |
| prairie | ERH93 | MCZ70291 | HU_MCZ | std/std | SRR13318747 |
| prairie | ERH94 | MCZ69534 | HU_MCZ | std/std | SRR13318736 |
| forest | ERH21 | MCZ69553 | HU_MCZ | std/inv | SRR13318612 |
| forest | ERH27 | MCZ69544 | HU_MCZ | inv/inv | SRR13318609 |
| forest | ERH36 | MCZ69566 | HU_MCZ | inv/inv | SRR13318608 |
| forest | ERH52 | MCZ69555 | HU_MCZ | inv/inv | SRR13318601 |
| forest | ERH19 | MCZ70244 | HU_MCZ | inv/inv | SRR13318613 |
| forest | ERH48 | MCZ69563 | HU_MCZ | inv/inv | SRR13318602 |
| forest | ERH26 | MCZ70248 | HU_MCZ | std/std | SRR13318610 |
| forest | ERH14 | MCZ70242 | HU_MCZ | inv/inv | SRR13318614 |
| forest | ERH25 | MCZ70247 | HU_MCZ | inv/inv | SRR13318611 |
| forest | ERH38 | NA | NA | inv/inv | SRR13318607 |
| forest | ERH42 | MCZ70254 | HU_MCZ | inv/inv | SRR13318606 |
| forest | ERH12 | MCZ70240 | HU_MCZ | inv/inv | SRR13318616 |
| forest | ERH43 | NA | NA | inv/inv | SRR13318605 |
| forest | ERH46 | MCZ69562 | HU_MCZ | inv/inv | SRR13318603 |
| forest | ERH53 | NA | NA | inv/inv | SRR13318600 |

